# Behavioral and transcriptomic changes following brain-specific loss of noradrenergic transmission

**DOI:** 10.1101/2023.01.12.523778

**Authors:** Elsa Isingrini, Chloé Guinaudie, Léa Perret, Elisa Guma, Victor Gorgievski, Ian D. Blum, Jessica Colby-Milley, Maryia Bairachnaya, Sébastien Mella, Antoine Adamantidis, Kay-Florian Storch, Bruno Giros

**Author notes:** Correspondence to BG.

## Abstract

Noradrenaline (NE) plays an integral role in shaping behavioral outcomes including anxiety/depression, fear, learning and memory, attention and shifting behavior, sleep-wake state, pain, and addiction. However, it is unclear whether dysregulation of NE release is a cause or a consequence of maladaptive orientations of these behaviors, many of which associated with psychiatric disorders.

To address this question, we used a unique genetic model in which the brain-specific Vesicular Monoamine Transporter-2 (VMAT2) gene expression was removed in NE-positive neurons disabling NE release in the entire brain. We engineered VMAT2 gene splicing and NE depletion by crossing floxed VMAT2 mice with mice expressing the Cre-recombinase under the DBH gene promotor. In this study, we performed a comprehensive behavioral and transcriptomic characterization of the VMAT2^DBHcre^ KO mice to evaluate the role of central NE in behavioral modulations.

We demonstrated that NE depletion induces anxiolytic and antidepressant-like effects, improves contextual fear memory, alters shifting behavior, decreases the locomotor response to amphetamine, and induces deeper sleep during the non-rapid eye movement (NREM) phase. In contrast, NE depletion did not affect spatial learning and memory, working memory, response to cocaine, and the architecture of the sleep-wake cycle. Finally, we used this model to identify genes that could be up- or down-regulated in the absence of NE release. We found an up-regulation of the Synaptic Vesicle Glycoprotein 2c (SV2c) gene expression in several brain regions, including the locus coeruleus (LC), and were able to validate this up-regulation as a marker of vulnerability to chronic social defeat.

## INTRODUCTION

Monoaminergic neurotransmitter systems, namely dopamine (DA), noradrenaline (NE), and serotonin (5HT), have similar general organization *and share common molecular properties*. Their cell bodies are localized in compact nuclei with extended projections to distant regions and they display multiple reciprocal connections [1]. These brainstem neuromodulatory systems extensively regulate behavioral states including mood, motivation, stress, arousal, vigilance, and attention and their role in neuropsychiatric disorders has been clearly established [2]. However, a better understanding of the relative contribution of each one of these systems in regulating behavioral state remains unclear and is complicated by their large *anatomical, pharmacological and functional overlaps*. At the molecular level, monoamines fluxes in the brain are primarily controlled by two types of transporters: (1) a single vesicular monoamine transporter-2 (VMAT2) [3–5] that actively transport all monoamines in presynaptic vesicles and (2) a specific plasma membrane transporters [6] that re-uptake the monoamines from the synaptic cleft.

Within the central nervous system (SNC), the NE system is comprised of clusters of neurons in the pons and medulla where the locus coeruleus (LC) is the largest nuclei. The LC receives inputs from a broad array of CNS structures and sends projections to nearly all regions of the brain and spinal cord to regulate cellular and local circuit activities [7–10]. The NE system is complex given that adrenoreceptors are ubiquitously present post-synaptically and pre-synaptically on neurons, glia, and blood vessels. According to the adrenoreceptor targeted, NE can have excitatory (α1 and β1, β2, β3) or inhibitory (α2 and in some cases β2) effects on neural activity by acting through Gα-mediated signaling. However, the NE modulatory effect is more complex than simply triggering an excitatory or inhibitory response. In numerous target areas, NE enhances the efficacy of synaptic transmission (either excitatory or inhibitory), while decreasing the spontaneous activity of the same neuron [11,12]. This complexity makes it difficult to predict how NE modulates the activity of brain circuits.

The NE facilitating effect on neural network functions is essential to optimize the behavioral outcome to extrinsic and intrinsic factors. In response to biological challenges, NE release has been observed in response to unexpected stimuli or conditioned to relevant stimuli [13,14]. Given the extensive projections of NE neurons throughout the CNS, NE response has been shown to modulate many behaviors. For example, NE projections to the thalamus and the cortex play a role in arousal and sensory processing [11,15–20]. NE signaling in the hippocampus (HPC) regulates synaptic plasticity [21–23] and together with projections to the amygdala influences memory consolidation [24,25]. In the prefrontal cortex (PFC), NE has been shown to regulate working memory, attention, and shifting behavior [26–29]. Moreover, NE also plays a role in regulatory networks of fear/anxiety and pain modulation [30,31]. This is consistent with the implication of NE neurons in controlling arousal and their activity across the sleep-wake cycle [14,32,33].

Our knowledge of how NE acts in the brain to modulate behavioral states is still incomplete. Multiple methodological approaches have been used to study the effects of NE system on brain function and behavior, including lesions (adrenalectomy and/or adrenergic denervation via surgical, immunological, or chemical lesions), pharmacological agents (stimulation or blockage of NE receptors via selective agonist or antagonists), and transgenic technology (knockout mice for NE synthesis enzyme dopamine β-hydroxylase (DBH) or tyrosine hydroxylase (TH)) combined with behavioral analysis and electrophysiological recording in target regions. All these techniques have contributed to the modern understanding of how NE participates in several regulatory networks in the brain, *albeit* there are limitations associated with each approach. Enzyme inhibition or chemical lesion only decreases 70% of NE levels [34–37]. The selective noradrenergic neurotoxin *N*-(2-chloroethyl)-*N*-2-bromobenzylamine (DSP-4) preferentially lesions neurons of the dorsal noradrenergic bundle originating from the LC, leaving the ventral noradrenergic bundle neurons relatively spared [38]. Moreover, given that DA is a precursor of NE synthesis, manipulating DA production without influencing NE concentrations is difficult (i.e. TH/DBH knockout mice, AMPT lesions) [39–44]. Other chemical lesions, including reserpine, deplete all monoamines or ablated co-transmitters as well as NE [45,46]. While these are important considerations, the main reason why nonspecific manipulation of NE concentration is challenging is that NE plays a key role in the peripheral nervous system, which is not the case for the other monoamines, DA and 5HT. This greatly complicates the use of drugs that interact with NE receptors or the genetic deletion of its synthesizing enzyme DBH [43] due to potentially severe side effects.

To overcome these limitations and investigate the exact role of central NE in behavioral states, we engineered a mouse model with selective and brain-specific NE depletion [47]. In this VMAT2^DBHcre^ mouse model, the vesicular monoamine transporter-2 (VMAT2) gene was specifically spliced-out in NE neurons by the Cre-recombinase expressed under the control of the DBH promoter. VMAT2 is essentially expressed in the CNS, whereas at the periphery NE and adrenaline (E) vesicle accumulation is under the control of the VMAT1 subtype [3,6]. This makes the VMAT2^DBHcre^ model able to fully preserve peripheral NE and E transmission. This model has been previously validated and the efficiency of VMAT2 splicing has been confirmed by in situ hybridization labeling in NE nuclei as well as by its consequences on NE metabolism [47]. The whole brain level of NE was dramatically decreased in KO mice compared to WT. To preclude any confounding effect on more complex behavior, we ensured that no alteration in the survival rate, growth, and motor abilities of animals was observed [47]. Here, we performed a detailed characterization of this mouse model in order to assess the role of CNS NE in regulating/modulating major brain networks associated with specific behavioral states. In parallel, we conducted a large transcriptomic analysis to identify genes whose expression was impacted by NE depletion in various brain regions, including both the LC and its projection areas, the ventral tegmental area (VTA), Raphe, Nucleus Accumbens (NAc), PFC and dentate gyrus DG.

## MATERIALS and METHODS

### Housing and breeding

Animal care and handling were performed according to the Canadian Council on Animal Care guidelines (CCAC; http://ccac.ca/en_/standards/guidelines) and approved by the Animal Care Committee of the Douglas Research Center.

The floxed VMAT2 mouse strain was produced at the Mouse Clinical Institute (Institut Clinique de la Souris, MCI/ICS, Illkirch, France). DBH-cre *[B6.FVB(Cg)-Tg(Dbh-cre)KH212Gsat/Mmucd*, stock number 036778-UCD] mice were obtained from the Mutant Mouse Regional Resource Center (MMRRC).

Heterozygous VMAT2 floxed mice (VMAT2^lox/+^) were crossed with heterozygous DBH^cre/+^ mice to obtain double heterozygote mice, which were then crossed to generate the WT (VMAT2^+/+^DBH^cre/+^) and KO (VMAT2^lox/lox^DBH^cre/+^) mice. Thereafter, VMAT2^lox/lox^DBH^cre/+^ KO mice are designated VMAT2^DBHcre^. All heterozygotes founders have more than 10 generations of backcross breeding on C57Bl/6J background. After weaning and sexing, males and females were separately housed in groups of 4-5 animals per cage and maintained under standard laboratory conditions: 22±1°C, 60% relative humidity, and a regular 12-12 h light-dark cycle (7:00-19:00 light period) with free access to food and water. The mice were used for behavioral screening at 2-4 months of age. When statistical analysis of sex revealed no differences, data for males and females were analyzed together. For corticosterone, nociception, circadian cycle, and sleep analysis, only male mice were used.

### Anxiety- and depression-like parameters

#### Elevated plus maze

The EPM was designed in a cross shape of four branching arms, with 2 opposing open arms (30 × 5 cm) and 2 opposing arms enclosed by a dark wall (30 × 5 ×11 cm). The arms radiated from a central platform (5 cm^2^), and the apparatus was 50 cm above the ground. The mice were placed in the central platform, facing an open arm, and were allowed to explore the maze for 5 min. An entry into the open or closed arm was defined when all 4 paws were inside the arm. The percentage of time spent in open arms was recorded as a behavioral parameter.

#### Novelty-suppressed feeding test

The test was conducted in an open field (45 × 45 × 45 cm) with a sawdust-covered floor under white illumination (40 W; ~2400 lux) positioned above the center of the open field. The mice were food-deprived for 24 hours prior testing. A single food pellet was placed on a round piece of white paper (12.5 cm diameter) at the center of the apparatus. Each mouse was placed in a corner of the open field with its head directed towards the wall, and the latency to eat was recorded up to a maximum testing period of 5 min. Immediately after, each animal was transferred to its home cage for 3 min, and the amount of food consumed was measured to assess changes in appetite as a confounding factor.

#### Marble burying test

Animals were placed in a cage filled with approximately 5 cm deep wood chip bedding, containing a regular pattern of 12 glass marbles on the surface (15 mm diameter, arranged in a 4 × 3 grid and spaced 4 cm apart). After 30 min, the number of marbles buried with bedding (2/3 their depth) was counted.

#### Forced swim test

The mice were dropped into an acrylic glass cylinder (height 25 cm, diameter 9 cm) filled with water at 21-23°C. Despair behavior, measured as immobility time, was scored during a 6-min test. Because little immobility is observed during the first 2 min of the test, immobility was only recorded during the remaining 4 min. Immobility was scored only when the mice ceased struggling and remained floating, motionless, and making only the movements necessary to keep their heads above water. Mice were intraperitoneally injected with either NaCl (0.9%), citalopram (diluted in NaCl 0.9%, 10 mg/kg), or reboxetine (diluted in NaCl 0.9%, 20 mg/kg), 30 minutes before the beginning of the test.

#### Sucrose preference test

Sucrose preference testing was carried out in the animal’s home cage in which 2 bottles were presented. The mice were habituated to the presence of two drinking water bottles for 2 days before being given the free choice of drinking either a 1% sucrose solution or regular water for a period of 4 days. Water and sucrose solution intake were measured daily by weighing the bottles, and the locations of the two bottles were switched daily to reduce any bias in side preference. Sucrose preference was calculated as a percentage of the weight of the sucrose bottle over the total weight of both the water and sucrose bottles; this was averaged over the 4 days of testing.

#### Restrain stress and saphenous blood collection

The conscious mouse was restrained in an uncapped 50 ml falcon tube that has air holes drilled into the closed end. The mouse’s nose was at the closed end of the tube with the back legs, rear, and tail of the animal exposed at the open end of the tube. The left hind leg was extended and fixed by firmly holding the fold of skin between the tail and thigh. To aid in the visualization of the saphenous vein, the hair was removed from the outer surface of the fixed leg with a small, sharp scalpel blade. Pinching the skin between the tail and thigh of the mouse restricts blood flow from the lower limb causing the saphenous vein to protrude. The shaved skin was wiped clean with 70% ethanol and then dried with a dry piece of gauze. In addition, a small amount of Vaseline was wiped onto the shaved skin to reduce clotting and prevent the blood from collecting in the remaining hair on the leg. A 25-gauge needle was held almost parallel to the saphenous vein and the vessel is punctured. Drops of blood were collected into a 0.5 ml EDTA-containing tube. After the first blood sampling (baseline), mice were kept in the falcon tube for 30 min before a second sampling (restrain stress). The mice then returned to their home cage for 90 min before the third sampling (return). Blood samples were centrifuged at 3000 rpm for 20 min and the plasma supernatant was removed and kept at - 80°C until analysis of corticosterone level. Blood samples were collected during the light phase between 9:00 and 12:00 hours.

#### Dexamethasone suppression test

To assess the HPA axis negative feedback to corticosterone, mice received a single intraperitoneal (i.p.) administration of either NaCl (0.9%) or of the glucocorticoid receptor agonist dexamethasone-phosphate (DEX-P, Sigma-Aldrich, St Louis, MO, USA) at a concentration of 0.5 mg/kg (dissolved in a 0.9% NaCl). Mice were sacrificed 2 hours after the injection and trunk blood was collected in 0.5 ml EDTA-containing tubes. Samples were centrifuged at 3000 rpm for 20 min and the plasma supernatant was removed and kept at −80°C until analysis of corticosterone level. Blood samples were collected during the light phase between 14:00 and 17:00 hours.

#### Corticosterone level measurement

Corticosterone plasma level was measured using commercially available immunoassay kits (Immunodiagnostic Systems) according to the manufacturer’s instructions. The intra-assay precision was ≤ 7%, the inter-assay precision was ≤ 9%, and the analytical sensitivity was 0.55 ng/mL.

### Learning and memory

#### Contextual and cued fear conditioning task

The fear conditioning apparatus consisted of two different contexts: (1) a black Plexiglas chamber measuring 20 cm wide, 18 cm high, and 28 cm deep with a stainless-steel grid floor (context A); and (2) a white Plexiglas chamber 20 cm wide, 18 cm high and 28 cm deep with black Plexiglas floor (Context B). The apparatus was connected to a shock generator. A 15W light bulb and a speaker to deliver the tone were located on the wall of the ceiling. The administration of the footshock (US) and auditory tone (CS) was controlled by computer. In the **conditioning session**, performed in context A, mice were placed in the test chamber and allowed to freely explore for a 2-min period. The adaptation period was followed by two pairings (2 min inter-trial interval (ITI)) of a 30-sec white noise period (CS; 80 dB) ending with a 2 sec 0.5 mA foot shock (US). Mice were removed from the conditioning chamber 30 sec after the termination of the last US. 24-hrs after the conditioning session, mice were tested for freezing response to the context or the tone. During the **contextual test**, performed in context A, mice were placed in the test chamber and their behavior scored for 3 min without the tone (CS) and the shock (US) presented. Three hours after the context test, mice were tested for freezing response to the CS. The **cued test**, performed in context B, consisted of a 2 min habituation period followed by 30 secs of white noise (CS; 80 dB). Throughout conditioning and the context and cued tests, mice behaviors were video recorded, and the continuous recording of the time spent freezing was scored. Freezing was defined as the absence of any visible movement but respiration.

#### Morris water maze

The water maze consisted of a circular stainless-steel pool (150 cm diameter, 29 cm height) filled with water maintained at 20–22°C and made opaque using a white aqueous emulsion. A circular escape platform (10 cm diameter) was submerged 1 cm below the water surface.

During the training of the **spatial version**, mice were trained to find the fixed position of the hidden platform, using extra-maze cues. Mice were released from pseudo-randomly assigned start locations and allowed to swim for up to 90 s when they were manually guided to the platform in the case of failures. Animals received one habituation trial on the first day and then two trials per day for 5 days (90 min inter-trial interval). On the seventh day, mice were given one additional training trial 90 min before the probe test, in which mice were allowed to swim for 60 s in the absence of the platform. Latencies to find the escape platform during training and the time spent in the active quadrant during the probe test were analyzed.

Mice were tested for a reversal training 24 hrs after the probe test, in which the same protocol is applied except that the hidden platform was moved to another location. Animals received two trials per day for 5 consecutive days (90 min inter-trial interval) and an additional training trial was performed on day 6 before the probe test. The time to find the escape platform during the training and time spent in the active quadrant during the probe test were analyzed.

During the **rapid place learning version** of the Morris water maze for working memory test, mice received 4 daily trials (2 min inter-trial interval) for 4 consecutive days. Every day, mice were trained to find the fixed position of the hidden platform using extra-maze cues, and the mice starting point was pseudo-randomly placed in different locations across the 4 trials. The position of the platform changed every day. Latencies to find the escape platform during the 4 days were analyzed.

### Executive functions

#### Attentional set-shifting task

The attentional set-shifting task is used as a measure of cognitive flexibility in mice. Prior testing, mice were individually housed and food restricted diet to maintain weight at 80-85% of their free-feeding body weight. The food restriction was maintained until the end of the testing. For habituation, mice were placed in the testing cage with two ramekins containing the food reward (chocolate chips) and filled with deep wood chip bedding. If a mouse failed to dig for the food reward after two successive days, it was excluded from the experiment and marked as a “failed to dig” in the experimental record. On the first day of testing, mice were given a **simple discrimination (SD)** test in which the two ramekins were filled with the same medium (wood chip bedding) and differed for the second relevant dimension (odor). For the **compound discrimination (CD)** test, a second medium was introduced, and the relevant dimension remained the odor. For the **reversal (CDR)**, the relevant dimension odor remained unchanged, but the previously correct stimulus was now incorrect. For the **intra-dimensional shift (ID)** and the **extra-dimensional shift (ED)**, new variants for both dimensions were used. In the ID shift, the relevant dimension was the same as before (odor) whereas in the ED shift the mouse had to shift attention to the previously irrelevant dimension (medium). On each day, mice were trained on a new task. As the paradigm was designed to examine the mice’s ability to shift from learning one task to another, the task learned on the previous day was repeated before introducing the new task. In each task, if the mouse started to dig in the incorrect ramekin, an error was recorded. The side of stimulus presentation varied pseudo-randomly. Tests continued until the mouse reached the criterion of six consecutive correct trials.

### Addiction-like behavior

#### Locomotor response

Locomotor activity was measured with an Omnitech digiscan activity monitor. To evaluate the effects of cocaine (5, 10, 20 mg/kg) or amphetamine (1, 3, 5 mg/kg) on locomotor behavior, the mice were first habituated for 1 hour to an open field chamber (21cm × 21cm) equipped with photocells and plexiglass walls and floors and then recorded for 2 hours after i.p. drug administration. The animal’s horizontal movements were measured in 5-min intervals for the 3-hour experiment.

#### Behavioral sensitization

To initiate sensitization, the mice were injected with cocaine (20 mg/kg, i.p.) or amphetamine (3 mg/kg, i.p.) for six consecutive days. After 1 day of withdrawal, on day 8, the mice were challenged with cocaine (20 mg/kg, i.p.) or amphetamine (3 mg/kg, i.p.) to test the expression of sensitization. The mice were tested for locomotor activity on days 1 and 8. The mice were habituated to the activity monitor cage (21 × 21 cm^2^) for 1 hour and then recorded for 2 hours after cocaine or amphetamine administration. Horizontal activity measured at 5 min intervals was compared between days 1 and 8 for the first 15 min of the cocaine challenge.

### Nociception

#### Tail immersion test

Each mouse was placed in a 50 ml Falcon plastic tube with the tail protruding from the opening side of the tube. The lower 5 cm section of mice tail was immersed in a bath of circulating water maintained at 48°C for a first session and then at 52°C. Latency to respond to the heat stimulus with vigorous flexion of the tail was recorded.

### Circadian Analysis

For all experiments, animals were individually housed with *ad libitum* access to food and water. For all genotypes and all conditions, seven continuous days of recording were used to generate parameter estimates and averages.

#### Running wheels

Animals were placed in cages equipped with running wheels in custom-built, light-controlled cabinets. Running wheel revolutions were recorded continuously (ClockLab, Actimetrics, Wilmette, IL). Actograms, displaying binned running wheel revolutions per 6 min (0.1 hr), and the associated Chi-squared period estimates were generated using ClockLab software as described [48].

#### Telemetry

Animals were housed in an isolated, climate-controlled room on a rack with both genotypes interleaved to avoid any confounds due to variable lighting on different shelves. Each mouse had *ad libitum* access to food and water in a standard cage placed atop energizer/receiver units (ER-4000, Starr Life Science Corp., Oakmont, PA). One week before data collection, electromagnetic induction powered telemetry probes (G2 E-mitter, Starr Life Science Corp.) were implanted intraperitoneally. Locomotion (counts) and core body temperature (°C), were collected in 6-min bins (0.1 hr) using Vitalview software (Starr Life Science Corp.). All telemetry data was exported into Clocklab for visualization and analysis [48].

### Sleep recording

#### Surgery

Under isoflurane anesthesia, mice were chronically implanted with stainless steel electroencephalogram/electromyogram (EEG/EMG) electrodes. Cortical electrodes were inserted into the skull bilaterally above the frontal (2mm lateral and anterior to bregma) and parietal cortices (2mm lateral to the midline at the midpoint between bregma and lambda, above the hippocampus). All cortical electrodes were fixed to the skull via SuperBond (Sun Medical Co., Shiga, Japan) and acrylic cement. Following surgery, animals were housed individually in their home cage for a recovery period of 15 days. Six to 10 days before polysomnographic recordings, the mice were transferred to a recording room and placed in individual recording cages where electrodes were connected to tether cables for habituation and electrophysiological recordings.

#### Polysomnographic recording and data acquisition

Twenty-four hour of baseline recordings were performed beginning at 20.00h (dark onset of the 12:12 light/dark cycle). EEG and EMG signals from electrodes were amplified (Grass Instruments, USA), digitized at a sampling rate of 512 Hz, and collected on a PC within the recording room using *VitalRecorder* (Kissei Comtec Co. Ltd, Japan). EEG and EMG signals were processed with low and high pass filters of 0.5–80 Hz and 20–40 Hz, respectively. Recordings were visually scored offline using *SleepSign* (Kissei Comtec Co. Ltd, Japan), using 5-second epoch window, as either wake, Non-rapid eye movement (NREM) sleep, or REM sleep, as previously described [49]. Briefly, the behavioral state was determined according to the following criteria:

-*Wake*: High-frequency low-amplitude EEG oscillations accompanied by constant EMG activity with phasic bursts.
-*NREM* sleep: Low frequency, high amplitude EEG oscillations with an increase in slow delta wave activity (0.5–4.5 Hz) and a loss of phasic muscle activity.
-*REM* sleep: High frequency, low amplitude EEG oscillations with typical regular theta rhythm (5-9 Hz) and a flat EMG.

#### Spectral power analysis

The cortical EEG signal of each behavioral state (wake, NREM, and REM sleep) was subjected to spectral analysis using Fourier transform to calculate the EEG power spectrum from 0.25-100 Hz. Power values were analyzed for the following frequency bands: delta (0.5-4.5 Hz), theta (5-9 Hz), and alpha (9-12 Hz). Spectral power values reported were normalized to the sum of all power values (V^2^) binned for every .025 Hz from 0.25-100 Hz.

### Quantitative in situ hybridization

3-month old VMAT2^DBHcre^ WT and KO mice brains were collected after decapitation and frozen in isopentane at −30°C. Brains were sliced in coronal sections (10 μm thick) using a cryostat (Leica CM3050S) and rinsed in 0.1 M PBS, SSC 1,0M and treated with 0.25% ethanol. [^35^S]-dATP oligonucleotides (**SV2c:** AS1: 5’-GAC TGT AGG ACC GCT GGG TAT ATT CAT CCT GGG CC −3’; AS2: 5’-GAC TGT AGG ACC GCT GGG TAT ATT CAT CCT GGG CC −3’; AS3: 5’-CTG CTG TAA CAG CTA GAG TGG CTG GCA GGC TGT CT −3’) were synthesized with terminal transferase (Amersham, Biosciences) to obtain a specific activity of 5×10^−8^ dpm/μg. Sections were covered with 70 μl of hybridization mix and 5×10^−5^ dpm of each labeled oligonucleotide and incubated overnight at 42°C in a humid chamber. Following washes and dehydration, slides were air-dried and exposed to a BAS-SR Fujifilm Imaging Plate for 5 days. The plates were scanned with a Fujifilm BioImaging Analyzer BAS-5000. Regions identification was based on Franklin and Paxinos Mouse Atlas [50].

### Microarray transcriptome

#### Tissue dissection, RNA isolation

3-month-old VMAT2^DBHcre^ KO and WT male mice were decapitated and brains flash frozen in isopentane −30°C and stored at −80°C. LC, VTA, Raphe, NAc, PFC and DG micropunch of 0.5 mm diameter were performed under −20°C cryostat. Tissue from three to five mice was pooled for WT (n=4) and KO (n=4) groups. RNA was isolated and purified with miRNAeasy micro kits from Qiagen (Cat#217084) and processed at Genome Quebec for RNA (Agilent - Mouse 60k) differential arrays.

#### Normalization, quality control, and filtering of microarrays data

Microarrays data were processed using standard quality control tools to obtain normalized, probeset-level expression data. The raw files were read, background corrected, and normalized using the Limma (v_3.30.13) bioconductor package [51,52] as follow: the raw files were read using the read.maimages function with the “agilent” option in the source parameter; the backgroundCorrect function was run with the “normexp” method and an offset of 18; then, we use the normalize between arrays with the “cyclicloess” method. Before fitting the linear model on the expression matrix, the low expressed probes and control probes were filtered out, for this, we compute the 95% percentile of the negative control probes on each array and kept the probes that were at least 10% brighter than the negative controls in at least four arrays since there were four replicates. Arrays quality control and sample distribution were examined using boxplots, hierarchical clustering of the Euclidean distance, and Principal Component Analysis.

#### Removing batch effects

For clustering and unsupervised analysis, batch effects were removed from the expression matrix using the removeBatchEffect function with the chip identity as a batch indicator.

For the differential expression analysis, the batch factors (i.e the chip identity) were included in the linear model, for this, the correlation between measurements was calculated with the duplicated Correlation function with the chip identity as input for the block argument and this correlation was used in the linear model fit as input of the correlation argument of the lmFit function.

#### Differential expression analysis

Each cerebral region was individually analyzed to identify probes showing significant differential expression (DEGs) between the wild type and the VMAT2^BDHcre^ KO samples. This analysis was performed using the linear model method implemented in the *Limma* R package. The basic statistic was the moderated t-statistic with a Benjamini and Hochberg’s multiple testing correction to control the false discovery rate (FDR) [53]. The gene symbols and Entrez Ids were added using the agilent annotation file, then duplicated genes were collapsed by keeping the one with the smallest adjusted p-value.

#### Gene set enrichment analysis

Tables for each region from the differential gene expression analysis were ordered by the t-statistics values allowing to rank the genes from the most upregulated to the most down-regulated. Gene set collections from the mouse version of the Molecular Signatures Database MSigDB v6 in R format were downloaded from Molecular Signatures Database (http://bioinf.wehi.edu.au/software/MSigDB/)

The gene collections were used to perform enrichment analysis using two complementary approaches: first, an over-representation analysis (ORA) (Khatri et al., 2012) on differentially expressed genes was performed using one-sided Fisher’s exact tests implemented in R [54] with a Benjamini and Hochberg’s multiple testing correction; second, a gene set enrichment analysis (GSEA) (functional scoring method (FSC) [55] was performed on the ranked list (see above) of genes using the runGSA function in piano R package [56].

#### Visual representation

Heatmaps were made using R package pheatmap (v_1.0.8), Venn diagrams were generated by gplots R package (v_ 3.0.1) and other visual representations (barplot and PCA biplot) were made using R package ggplot2 (v_2.2.1).

#### R Session info

All analyses were performed using R (R Core Team, 2017) version 3.3.2 (2016-10-31), running on OS X El Capitan 10.11.6 on x86_64-apple-darwin13.4.0 (64-bit) platform.

### Statistical analysis

Statistical analyses were performed using Statistica software. The Shapiro-Wilk test was used to check whether the sample distribution is normal and the Levene’s test was used to evaluate the homogeneity of variances. The results are expressed as mean ± SEM (standard error of the mean). According to the experimental design, multiple group comparisons were performed using a two-way factorial ANOVA or a one-way repeated measure ANOVA. The between-subject factors were: genotype (WT *vs* KO), treatment (FST: NaCl *vs* Citalopram *vs* reboxetine), tone (Fear conditioning: No tone *vs* Tone), temperature (Tail immersion test: 48°C *vs* 52°C), concentration (Acute locomotor drugs response: cocaine 5 mg/kg *vs* 10 mg/kg *vs* 20 mg/kg; amphetamine 1 mg/kg *vs* 3 mg/kg *vs* 5 mg/kg), cycle (Circadian analysis: L:D *vs* D:D), state (Wake *vs* NREM *vs* REM). The within-subject factors were: Day (Sucrose preference test and drug sensitization), Time (Corticosterone level analysis), Session (MWM training), Test (Set shifting), Hour (sleep recording from 20h to 8h), and Frequency (Sleep recording from 0 to 12 Hz). A Fisher LSD *post-hoc* test for pair-wise comparisons was then applied when appropriate. For the comparison of two independent groups (VMAT2^DBHcre^ WT *vs* KO) in the EPM, NSF, marble burying, dexamethasone suppression test, fear contextual test, the MWM probe test, and the telemetric body temperature, we used a two-sided t-test by groups. Optic density was quantified using MCID for SV2c mRNA labeling in the LC, VTA, and NAc. A two-sided t-test by groups was used for comparison between WT and KO groups and a one-way ANOVA followed by a Fisher LSD *post-hoc* test was used to compare SV2c mRNA density between control, susceptible and resilient mice to chronic social defeat stress. P<0.05 was selected to reflect statistically significant differences between groups.

## RESULTS

### Role of central NE system in anxiety and depression

To determine NE implication in anxiety-like behavior, we used the elevated plus maze, the novelty suppressed feeding test (NSF), and the marble burying test. While VMAT2^DBHcre^ KO mice showed a tendency to spend more time in the open arms in the elevated plus maze compared to WT, this effect was not statistically significant (t_56_=−1.5, p=0.14, *ns*; Figure 1A). However, significant effects were found in the NSF test and the marble burying test. In the NSF test, the latency to chew the food pellet placed in the center of an open field was significantly decreased in VMAT2^DBHcre^ KO mice compared to their WT littermate (t_50_=3.70, p=0.0005; Figure 1B). However, the amount of food consumed in the home cage at the end of the test was identical between genotypes (t_50_=−1.51, p=0.14, *ns*, data not shown), indicating that change in appetite was not a confounding effect on the observed decrease latency in VMAT2^DBHcre^ KO mice. In the marble burying test, VMAT2^DBHcre^ KO mice buried significantly less marbles than WT over 30 minutes (t_33_=2.07, p=0.047, Figure 1C). These results suggest that NE may play a critical role in mediating anxiety-like behavior with an “anxiolytic-like” effect of NE depletion in VMAT2^DBHcre^ mice.

**Figure 1:**
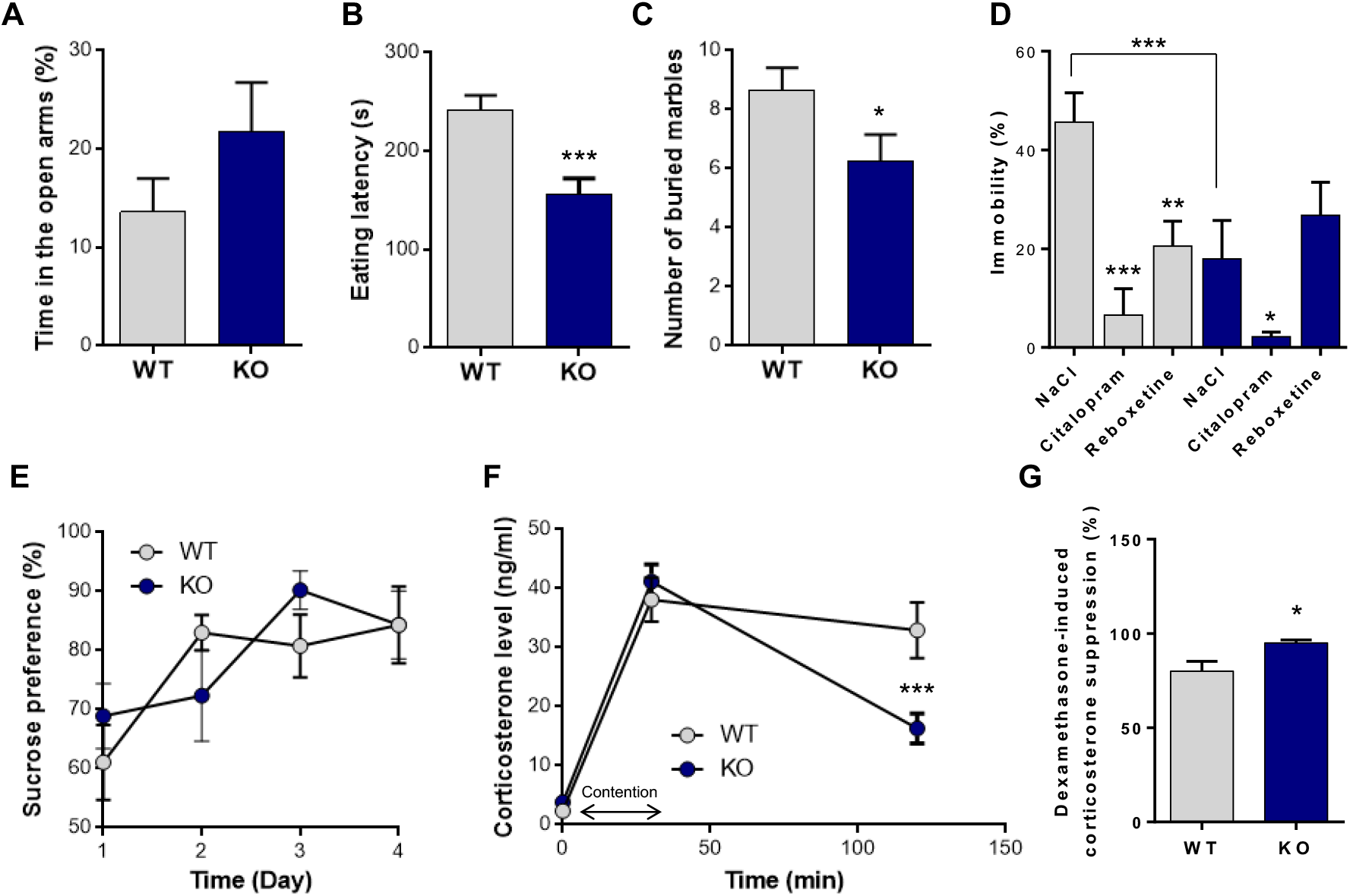
Anxiety and depression-like behavior in VMAT2^DBHcre^ mice. **A.** Percentage of time spent in the open arms of the elevated plus maze in VMAT2^DBHcre^WT (n=29) and KO (n=29) mice (t_56_=−1.5, p=0.14, *ns*). **B.** Latency (s) to eat the food pellet placed at the center of the open field in the novelty-suppressed feeding test in VMAT2^DBHcre^ WT (n=25) and KO (n=27) mice (t_50_=3.70, ***p=0.0005). **C.** Number of buried marbles with bedding during 30 minutes in VMAT2^DBHcre^ WT (n=18) and KO (n=17) mice (t_33_=2.07, *p=0.047). **D.** Percentage of immobility time in the forced swim test in VMAT2^DBHcre^ WT and KO mice injected intraperitoneally with NaCl (0.9%; WT n=8, KO n=8), citalopram (10 mg/kg; WT n=6, KO n=8) and reboxetine (20 mg/kg; WT n=8, KO n=6; Genotype × Treatment: F_(2,38)_=4.73, p=0.015; *post-hoc* test: compared to the NaCl group of the same genotype *p=0.048 **p=0.002 ***p=0.00004, compared to WT group with the same treatment ***p=0.0009). **E.** Percentage of sucrose preference over drinking water during 4 consecutive days in VMAT2^DBHcre^ WT (n=10) and KO (n=9) mice (Genotype × Day: F_(3,48)_=0.81, p=0.49, *ns*). **F.** Plasma corticosterone level (ng/ml) at baseline, after 30 minutes of restrain stress and 90 minutes after the end of the stress in VMAT2^DBHcre^ WT (n=18) and KO (n=17) mice (Genotype × Time: F_(2,66)_=7.63, p=0.001; *post-hoc test:* compared to the WT group at the same time ***p=0.00017). **G.** Percentage of dexamethasone-induced suppression of plasma corticosterone level in VMAT2^DBHcre^ WT (n=5) and KO (n=5) (t_8_=2.72, *p=0.026).

We next investigated the effect of NE depletion on depression-like behavior and associated physiological changes. In the forced swim test, acute treatment with citalopram (selective serotonin reuptake inhibitor, SSRI) and reboxetine (selective noradrenaline reuptake inhibitor, SNRI) induced a significant decreased in immobility in WT animals (F_(2,38)_=4.73, p=0.015; *post-hoc*: citalopram p=0.00004, reboxetine p=0.002; Figure 1D). In contrast, while citalopram decreased immobility (*post-hoc*: p=0.048), reboxetine had no effect in VMAT2^DBHcre^ KO mice (*post-hoc* test: p=0.30, *ns*), consistent with the fact that SNRI reboxetine is unable to induce behavioral changes in the absence of NE release in KO. Moreover, KO mice had a significantly decreased immobility as compared to their WT littermate, (*post-hoc* test: p=0.0009, Figure 1D) indicating that NE depletion may have antidepressant-like effects in this test. Finally, WT and KO mice showed no differences in sucrose preference measured over 4 consecutive days (F_(3,48)_=0.81, p=0.49, *ns*, Figure 1E). Physiological changes associated with depression were tested by measuring plasma corticosterone levels and quantifying the HPA axis negative feedback. The corticosterone level of KO mice in the basal condition and 30 minutes following the restrain stress was unchanged compared to WT (F_(2,66)_=7.63, p=0.001; *post-hoc* test: basal: p=0.72, ns, restrain stress p=0.47, *ns*). However, the decrease of corticosterone 90 minutes after the end of the stress exposure was faster in KO than in WT mice (*post-hoc* test: p=0.00017, Figure 1F), which may reflect improved HPA axis negative feedback to control corticosterone release in KO mice. To test this hypothesis, we performed the dexamethasone suppression test, which allows assessing the integrity of the HPA axis negative feedback. We observed that the percentage of dexamethasone-induced suppression of plasma corticosterone level was increased in KO mice compared with WT (t_8_=2.72, p=0.026, Figure 1G), indicating more efficient negative feedback in KO mice with NE depletion. Overall, these suggest that central NE depletion may have behavioral and physiological antidepressant-like effects in basal conditions and in response to acute stress.

### NE and fear conditioning

Considering that VMAT2^DBHcre^ KO mice were less anxious on our behavioral measures and showed less vulnerability to acute stress, we tested the VMAT2^DBHcre^ mice for emotional memory with two components, contextual recall and cued memory (Figure 2A). In the fear conditioning test, while no alteration was found in the cued test (F_(1, 13)_=0.053, p=0.82, *ns*), VMAT2^DBHcre^-KO mice exhibited a significant increase in freezing behavior during the contextual recall test (t_13_=2.86, p=0.013), indicating a specific impairment in contextual emotional memory. This effect was not due to altered pain sensitivity threshold as nociceptive reaction in VMAT2^DBHcre^ mice was similar to that of WT mice using the tail immersion test at both 48°C and 52°C (Figure 2B).

**Figure 2:**
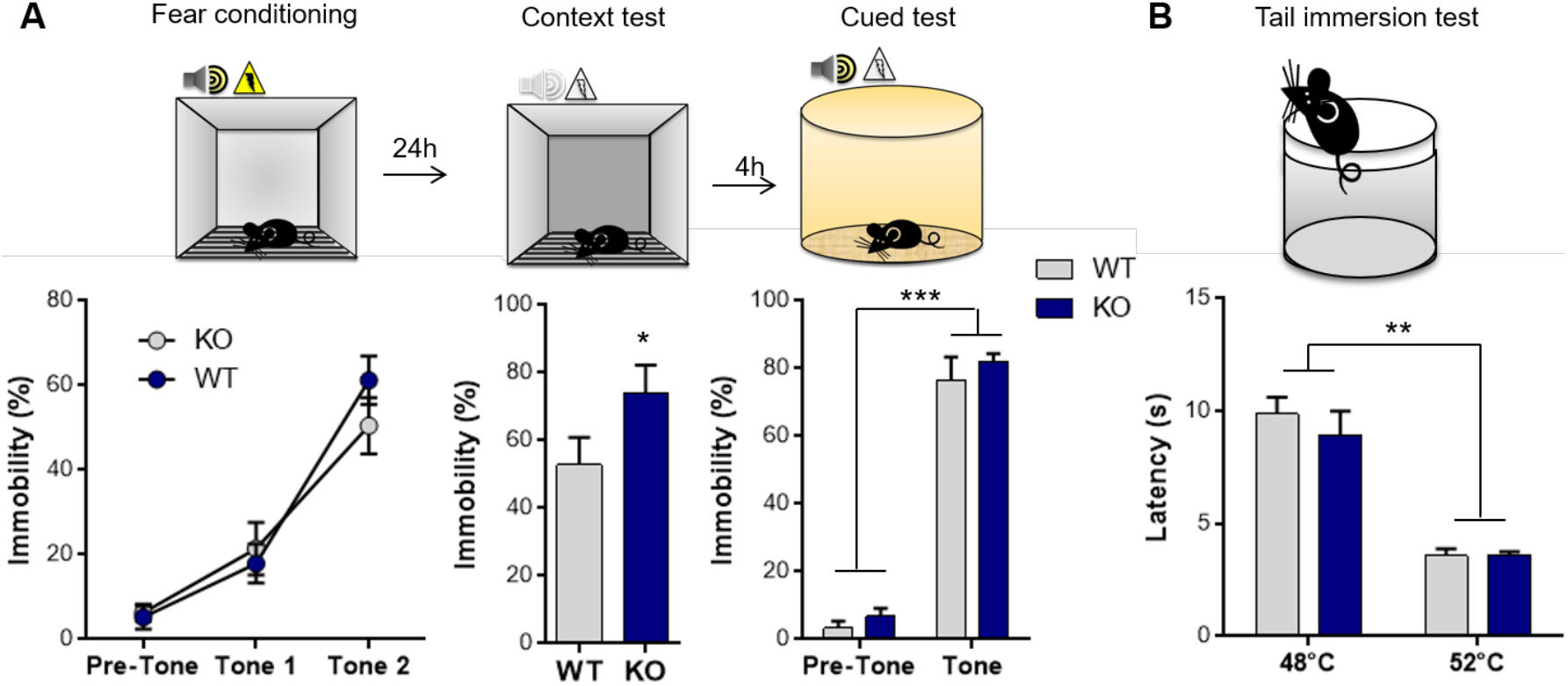
Emotional memory and nociception in VMAT2^DBHcre^ mice. **A.** Percentage of immobility time during the 2-min adaptation period and the two pairings (2 min ITI) of a 30-sec tone (80 dB) ending with a 2 sec 0.5 mA foot shock in VMAT2^DBHcre^ WT (n=7) and KO (n=8) mice (Genotype: F_(1,13)_=0.17, p=0.69; Tone: F_(2,26)_=76.20, ***p<0.001; Genotype × Tone: F_(2,26)_=1.67, p=0.21; left). Percentage of immobility during the 3 min of the contextual fear memory test performed 24h after the fear conditioning (t_13_=2.86, *p=0.013; middle). Percentage of immobility during the 2-min habituation and the 30-sec tone (80dB) in the cued memory test performed 24h after the fear conditioning (Genotype: F_(1,13)_=1, p=0.33; Tone: F_(1,13)_=365.25, p<0.001; Genotype × Tone: F_(1, 13)_=0.053, p=0.82; right). **B.** Latency (s) to respond to the heat stimulus with vigorous flexion of the tail when the water temperature is maintained at 48°C for a first session and then at 52°C in VMAT2^DBHcre^ WT (n=10) and KO (n=9) mice (Genotype: F_(1,17)_=0.44, p=0.52; Temperature: F_(1,17)_=92.91, ***p<0.001; Genotype × Temperature: F_(1,17)_=0.57, p=0.46).

### NE and memory

We then evaluated the cognitive performance of the animals using a different version of the Morris water maze paradigm (MWM; see Methods). In the long-term memory recall of the spatial version of the MWM, no deficit was observed in the VMAT2^DBHcre^ KO mice in comparison to WT. The latency to reach the hidden platform using spatial cues along the training sessions (F_(6.108)_=1.25, p=0.29, *ns*) and the time spent in the active quadrant during the memory test (t_18_=−0.34, p=0.74, *ns*) were similar between WT and KO mice (Figure 3A). In the rapid-place learning version of the MWM test, no working memory deficit was found in VMAT2^DBHcre^ KO mice compared to WT. Over the 3 days of training, the latency to find the hidden platform using spatial cues was similar between genotypes (Day 1: F_(3,54)_=0.55, p=0.65, *ns*; Day 2: F_(3,54)_=0.90, p=0.45, *ns*; Day 3: F_(3,54)_=0.67, p=0.57, *ns*) (Figure 3B).

**Figure 3:**
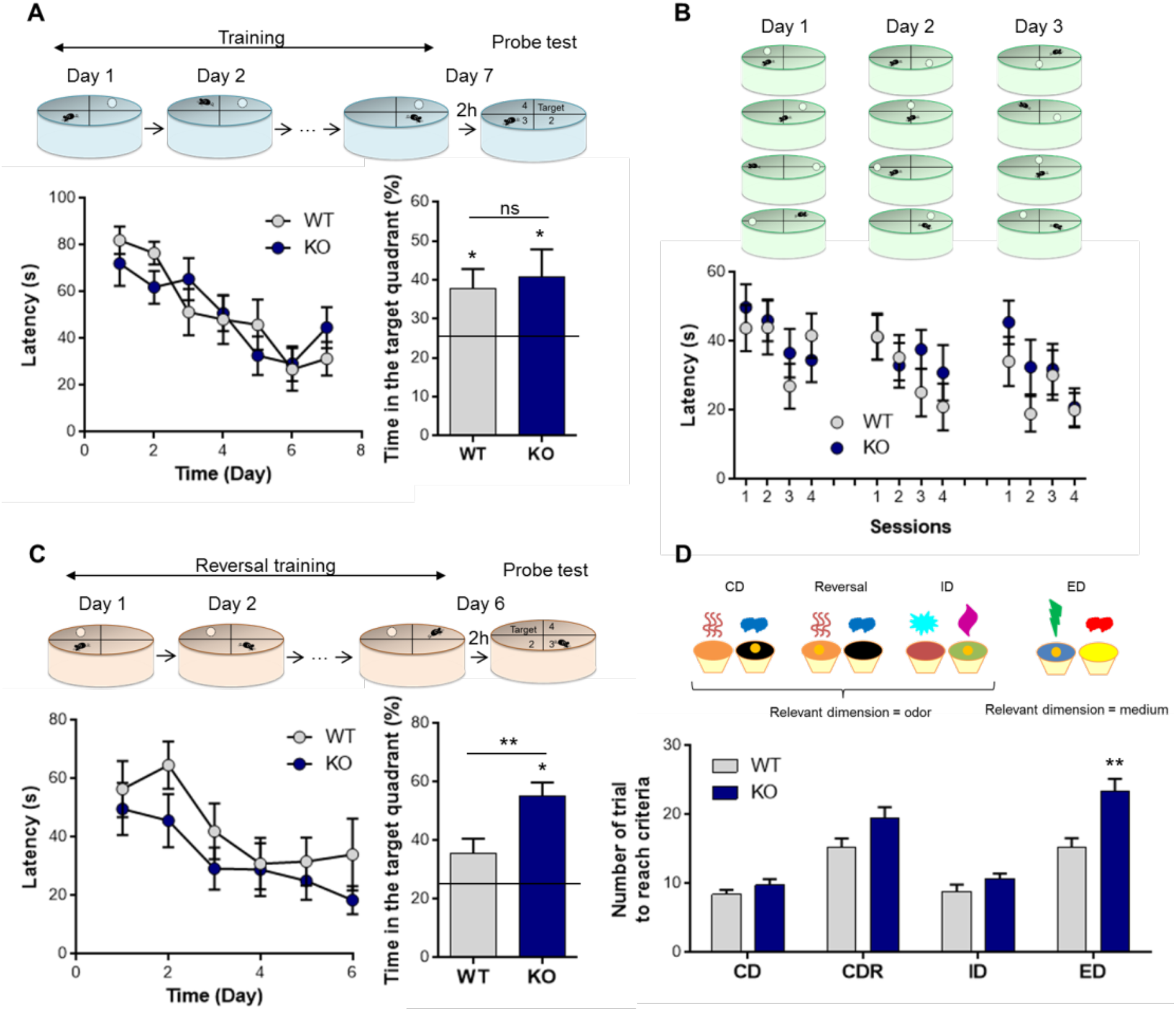
Learning, memory and adaptative behavior in VMAT2^DBHcre^ mice. **A.** Mean escape latency (s) in the spatial hidden-platform version of the Morris water maze in VMAT2^DBHcre^ WT and KO (n=10 per group, Left) (Genotype: F_(1,18)_=0.012, p=0.91; Session: F_(6,108)_=10.86, p<0.001; Genotype × Session: F_(6,108)_=1.25, p=0.29). Percentage of time spent in the active quadrant during the probe test of the MWM in each genotype (WT vs KO: t_18_=−0.34, p=0.74 (ns); Difference to 25%: WT t_6_=2.45, *p=0.037; KO tg=2.17, *p=0.05). **B.** Escape latency (s) to retrieve the hidden platform during 4 sessions per day for 3 days in the rapid place learning version of the Morris water maze for working memory test in VMAT2^DBHcre^ WT (n=10) and KO (n=10) mice (Day 1: Genotype F_(1,18)_=0.36, p=0.56; Session F_(3,54)_=2.02, p=0.12; Genotype × Session F_(3,54)_=0.55, p=0.65; Day 2: Genotype F_(1,18)_=0.53, p=0.47; Session F_(3,54)_=2.75, p=0.051; Genotype × Session F_(3,54)_=0.90, p=0.45. Day 3: Genotype F_(1,18)_=1.24, p=0.28; Session F_(3,54)_=4.33, p=0.008; Genotype × Session F_(3,54)_=0.67, p=0.57). **C.** Mean escape latency (s) in the reversal training of the spatial hidden-platform version of the Morris water maze in VMAT2^DBHcre^ WT and KO (n=10 per group, Left)(Genotype: F_(1,16)_=1.96, p=0.18; Sessions: F_(5,90)_=5.69, p=0.0001; Genotype × Session: F_(5,90)_=0.35, p=0.88). Percentage of time spent in the active quadrant during the probe test in each genotype (WT vs KO: t_18_=−2.77, p=0.013; Difference to 25%: WT t_9_=2.06, p=0.069; KO t_9_=6.22, ***p=0.00015).**D.** Number of trials required to found the food reward in order to reach the criteria of 6 consecutive correct trials during the compound discrimination test (CD), the reversal test, the intra-dimensional (ID) shift and the extra-dimensional (ED) shift in VMAT2^DBHcre^ WT (n=15) and KO (n=17) mice (Genotype: F_(1,30)_=18.22, p=0.00018, Test: F_(3,90)_=40.76, p<0.0001, Genotype × Test: F_(3,90)_=3.51, p=0.018; *post-hoc* test *p=0.016, ***p<0.001).

### NE depletion and behavioral flexibility

To investigate adaptative behavior (executive function), we used the reversal version of the MWM and the attentional set-shifting task. In the reversal version of the MWM, the latency to find the platform using spatial cues during the training (F_(6,108)_=1.25, p=0.29, ns) were identical between genotypes. However, KO mice spend more time in the active quadrant than WT mice during the probe test (t_18_=2.77, p=0.0.013) (Figure 3C). In contrast, VMAT2^DBHcre^ KO mice required an increased number of trials to reach criteria compared with WT in both the reversal (F_(3,90_)=3.51, p=0.018; *post-hoc* test p=0.016) and in the extra-dimensional shift stage (*post-hoc* test p<0.001) of the attentional set-shifting test (Figure 3D).

### Role of NE depletion in drugs locomotor response

Following an acute cocaine injection, KO and WT mice had a similar locomotor response regardless of the concentration of cocaine used (F_(2,30)_=1.01, p=0.38, *ns*; Figure 4A). In contrast, acute administration of amphetamine led to hyperlocomotion in WT but not in the VMAT2^DBHcre^ KO, at all concentrations tested (F_(1,35)_=6.87, p=0.013; Figure 4C). To measure motor characteristics following chronic drug treatment, mice were subjected to a cocaine or amphetamine sensitization paradigm. Cocaine (i.p. 20 mg/kg; Figure 4B) or amphetamine (i.p. 3mg/kg; Figure 4D) was injected daily over 6 consecutive days. Two days after the last injection, the mice were challenged with the same dose of cocaine or amphetamine respectively. This paradigm consistently resulted in a significant heightened locomotor response to the challenge dose of cocaine or amphetamine in both the WT and the KO animals (Days effect: cocaine F_(1,18)_=54.04, p<0.001; amphetamine Day: F_(1,16)_=55.92, p<0.001), however, no difference was observed between genotypes (Cocaine: F_(1,18)_=0.64, p=0.43, *ns*; Amphetamine F_(1,16)_=0.0081, p=0.93, *ns*).

**Figure 4:**
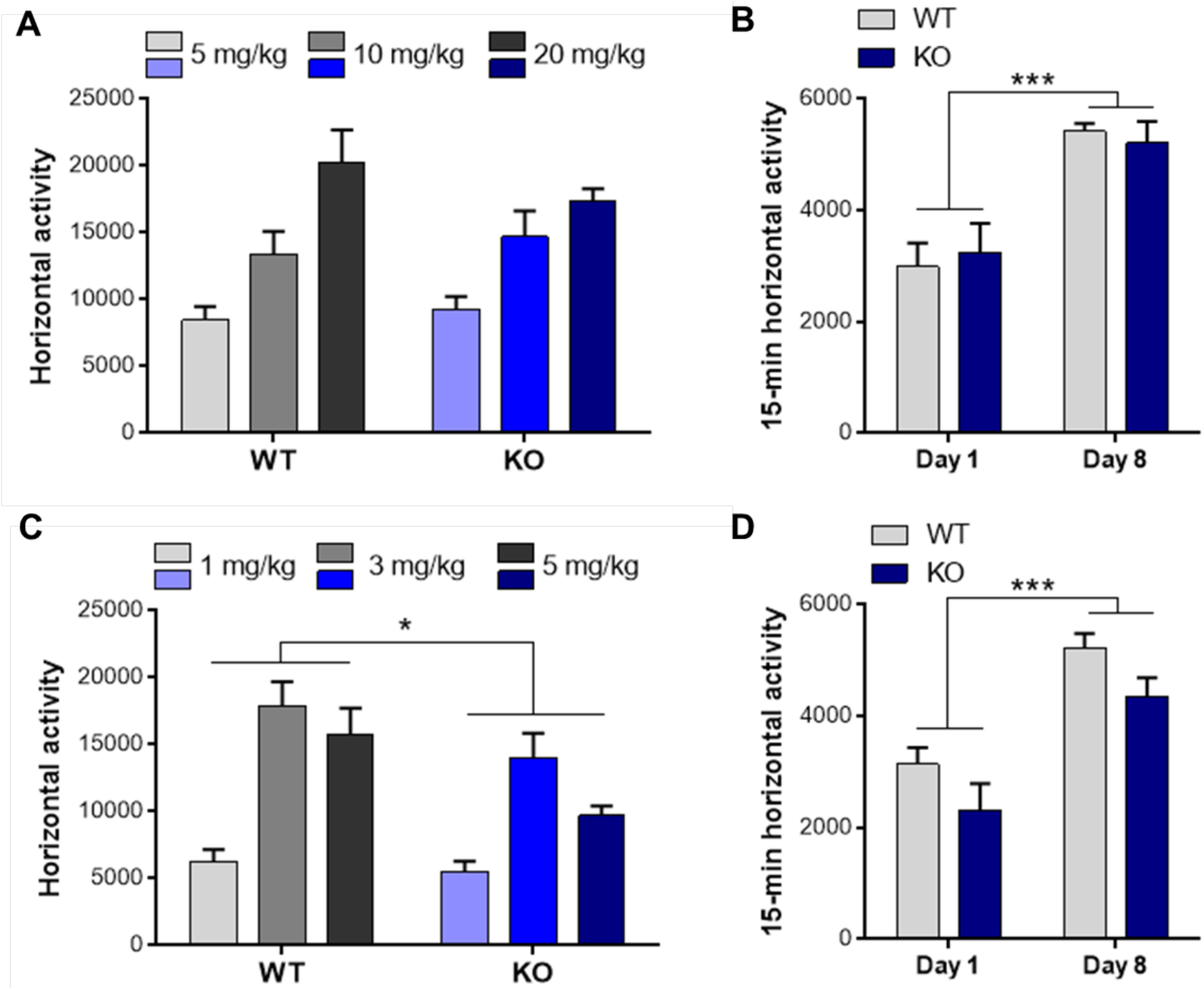
Addiction-like behavior in VMAT2^DBHcre^ mice. **A.** Acute locomotor response (horizontal activity) to cocaine during 2h following i.p. injection of 5, 10 or 20 mg/kg in the WT and KO mice of the VMAT2^DBHcre^ (n-6 per group; Genotype: F_(1,30)_=0.04, p=0.84; Concentration: F_(2,30)_=19.6, p<0.001; Genotype × Concentration: F_(2,30)_=1.01, p=0.38). **B.** Locomotor sensitization after repeated i.p. administration of 20 mg/kg cocaine over 6 days followed by 48 hours of withdrawal in the WT and KO mice of the VMAT2^DBHcre^ (n=10 per group; Genotype: F_(1,18)_=0.003, p=0.95; Day: F_(1,18)_=54.04, ***p<0.001, Genotype × Day: F_(1,18)_=0.64, p=0.43). **C.** Acute locomotor response (horizontal activity) to amphetamine during 1h following i.p. injection of 1, 3 or 5 mg/kg in the WT (1 mg/kg n=5, 3 mg/kg n=9, 5 mg/kg n=6) and KO (1 mg/kg n=6, 3 mg/kg n=9, 5 mg/kg n=6) mice of the VMAT2^DBHcre^ (Genotype: F_(1,35)_=6.87, *p=0.013; Concentration: F_(2,35)_=19.2, p<0.001; Genotype × Concentration: F_(2,35)_=1.13, p=0.33). **D.** Locomotor sensitization after repeated i.p. administration of 3 mg/kg amphetamine over 6 days followed by 48 hours of withdrawal in the WT and KO mice of the VMAT2^DBHcre^ (n=9 per group; Genotype: F_(1,16)_=4.11, p=0.06; Day: F_(1,16)_=55.92, ***p<0.001, Genotype × Day: F_(1,16)_=0.0081, p=0.93).

### NE and circadian rhythms

We studied circadian behavior of the VMAT2^DBHcre^ mice by monitoring their activity in wheel-running cages or by telemetry. Mice were entrained to a 12h/12h light/dark (L:D) cycle for 7 days and then transferred to constant darkness (D:D). Regardless of the mode of locomotor activity recording (running wheels/telemetry), the circadian locomotor periods of KO and WT mice were indifferent during both L:D and in constant darkness (D:D) (Running wheel: F(1,16)=0.22, p=0.64, *ns*; Telemetry: F(1,10)=0.0, p=1, *ns*; Figure 5A). The 7-day average body temperature measured by telemetry during L:D was similar between WT and KO mice (t_10_=−0.18, p=0.86, *ns*; Figure 5B), a result confirming that VMAT2 removal in DBH positive neurons does not affect peripheral NE release controlling the autonomic nervous system. However, total daily activity derived from wheel running was significantly decreased in KO when compared to WT regardless of lighting condition (F_(1,16)_=11.39, p=0.0039; Figure 5C-D), while total daily telemetry-derived activities were similar between genotypes during both L:D and D:D (F_(1,10)_=0.0071, p=0.93, *ns*; Figure 5E-F).

**Figure 5:**
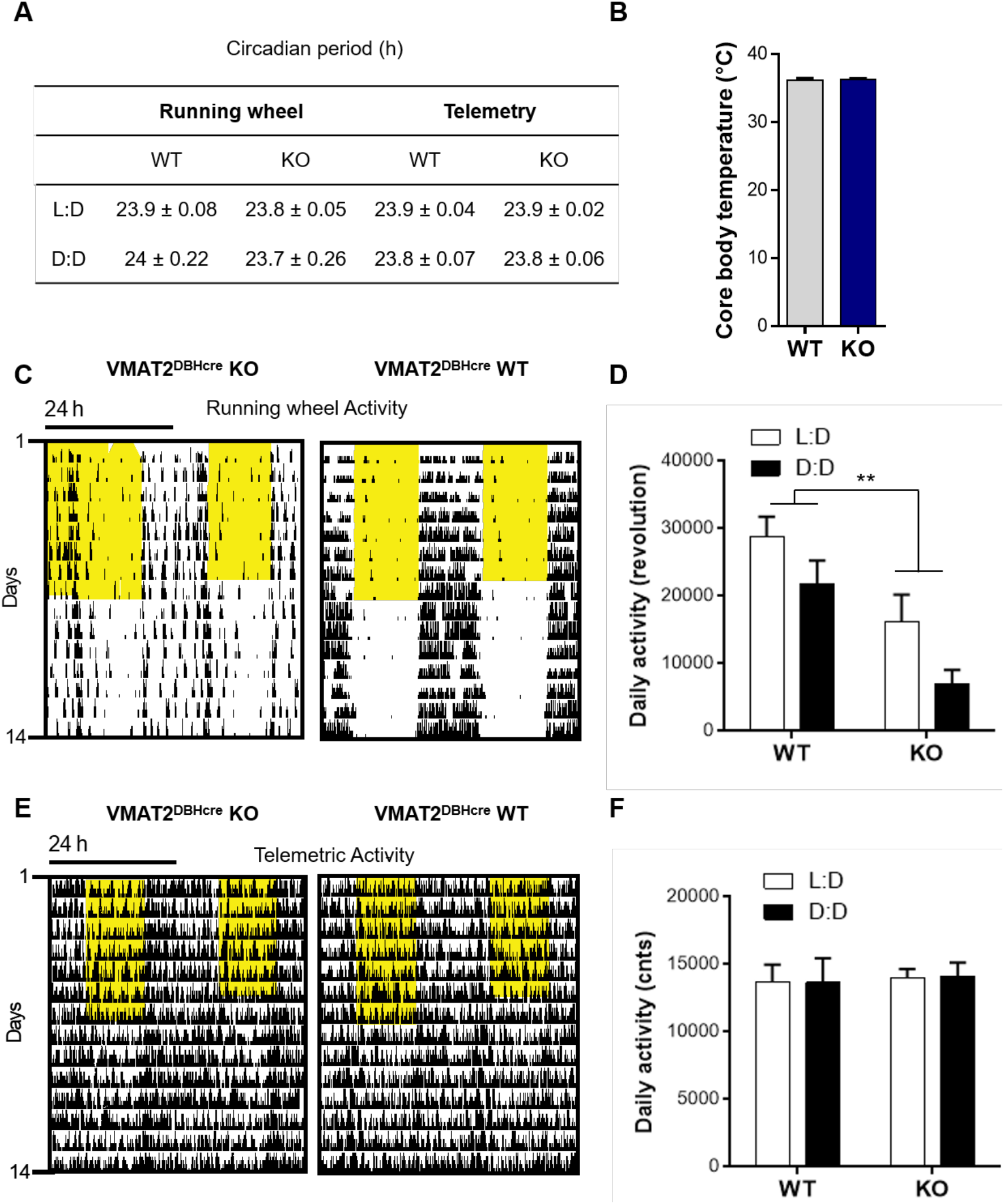
Circadian rhythmicity in VMAT2^DBHcre^ mice. **A.** Average circadian locomotor period derived from Chi-Squared periodogram analysis of 7-day time-spans during light/dark (L:D) cycles and constant darkness (D:D) derived from homecage running wheel activity (Genotype: F(1,16)=0.77, p=0.39, Cycle: F(1,16)=0.01, p=0.91, Genotype × Cycle: F(1,16)=0.22, p=0.64) or telemetry implants (Genotype: F(1,10)=0.0, p=1, Cycle: F(1,10)=11.6, **p=0.006, Genotype x Cycle: F(1,10)=0.0, p=1). **B.** Average core body temperature (°C) in WT (n=6) and KO (n=6) VMAT2^DBHcre^ mice measured by telemetry over 7 days (t_10_=−0.18, p=0.86). **C.** Representative double-plotted actograms of running wheel activity in VMAT2^DBHcre^ WT (right) and KO (left) mice. **D.** Mean running wheel activity over 7 days during L:D and D:D in VMAT2^DBHcre^ WT (n=9) and KO (n=9) mice (Genotype: F_(1,16)_=11.39, **p=0.0039; Cycle: F_(1,16)_=17.43, p<0.001; Genotype × Cycle: F_(1,16)_=0.32, p=0.58). **E.** Representative double-plotted actograms of telemetric activity in VMAT2^DBHcre^ WT (right) and KO (left) mice. **F.** Mean telemetric activity over 7 days of L:D and D:D in VMAT2^DBHcre^ WT (n=6) and KO (n=6) mice (Genotype: F_(1,10_)=0.055, p=0.82; Cycle: F_(1,0)_=0.00, p=0.99; Genotype × Cycle: F_(1,10)_=0.0071, p=0.93).

### NE control of sleep and arousal

Since NE is one of the major arousals modulators in the brain, we characterized the sleep-wake cycle of the VMAT2^DBHcre^ KO mice. We observed that the overall duration of wake, NREM, and REM sleep were unchanged between the two genotypes despite a decreased percentage of time spent awake in KO mice at 1h and 4h during the dark phase (F_(23,138)_=1.73, p=0.028; *post-hoc* test *p< 0,036, **p< 0,003), concomitant with an increase of NREM sleep (F_(23,138)_=1.62, p=0.047; *post-hoc* test *p< 0,043, **p< 0,0037) and no change in REM sleep (Figure 6A). Furthermore, the number of waking, NREM, and REM sleep bouts during both the light phase (F_(2,27)_=0.39, p=0.68) and the dark phase (F_(1,10)_=0.042, p=0.96) were similar between WT and KO mice (Figure 6B), as well as the duration of bouts ((Left; Genotype: F_(1,27)_=1.33, p=0.26; State: F_(2,27)_=99.72, p<0.001; Genotype × State: F_(2,27)_=0.40, p=0.68) and the dark cycle (Right; Genotype: F_(1,27)_=0.27, p=0.61; State: F_(2,27)_=34.03, p<0.001; Genotype × State: F_(2,27)_=0.043, p=0.96); Figure 6C). Upon examination of the power spectrum of the cortical EEG during NREM episodes, we found a significant increase in slow-wave activity (0.5-4.5 Hz) in KO mice compared to WT, both during the light (Curve: F(48,384)=2.5, p<0.001; *post-hoc* test *p< 0.05, **p<0.01, ***p<0.001; Histo: F_(2,24)_=9.31, p=0.001; *post-hoc* test ***p< 0,00032) and the dark phases (Curve: F_(48,384)_=3.64, p<0.001; *post-hoc* test *p< 0.05, **p<0.01, ***p<0.001; Histo: F_(2,24)_=4.70, p=0.019; *post-hoc* test *p< 0,015; Figure 7A), indicative of a deeper sleep intensity (i.e., higher slow waves amplitude) during NREM in NE-depleted mice. Minor, yet significant, changes in the power spectrum of cortical EEG analysis included a decrease in slow wave activity (Delta: 0.5-4.5 Hz) during the dark phase (F_(2,24)_=9.2, p=0.001, *post-hoc* test ***p=0.0005; Figure 7B) for awake episodes and an increase in theta rhythms (5-9 Hz; F_(2,24)_=4.37, p=0.024, *post-hoc* test ***p=0.0086; Figure 7C) restricted to the light phase of REM sleep episodes.

**Figure 6:**
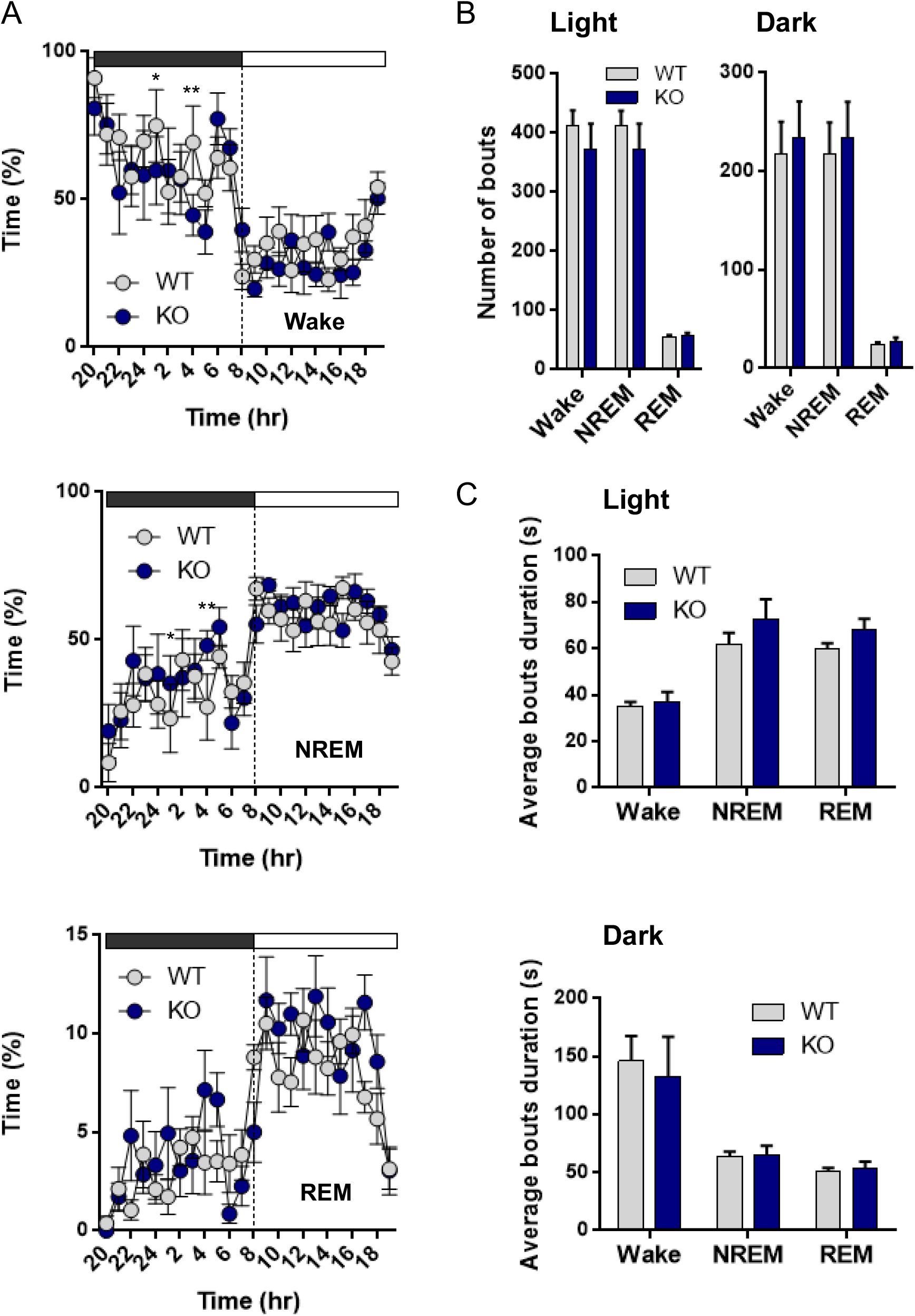
Sleep architecture analysis in VMAT2^DBHcre^ mice. **A**. Hourly percentage of time over a 24h period during the dark (grey) and the light cycle (white) spent in wake (Top; Genotype: F_(1,138)_=0.46, p=0.52; Hour: F_(23,138)_=7.03, p<0.001; Genotype × Hour: F_(23,138)_=1.73, p=0.028; *post-hoc* test *p< 0,036, **p< 0,003), non-rapid eye movement (NREM, middle; Genotype: F_(1,138)_=0.35, p=0.57; Hour: F_(23,138)_=6.34, p<0.001; Genotype × Hour : F_(23,138)_=1.62, p=0.047; *post-hoc* test *p< 0,043, **p< 0,0037) and REM (Bottom; Genotype: F_(1,138)_=2.1, p=0.2; Hour: F_(23,138)_=7.01, p<0.001; Genotype × Hour: F_(23,138)_=1.56, p=0.062) in WT (n=6) and KO (n=5) VMAT2^DBHcre^ mice. **B**. Number of bouts spent in wake, NREM and REM during the light (Left; Genotype: F_(1,27)_=1.33, p=0.26; State: F_(2,27)_=99.72, p<0.001; Genotype × State: F_(2,27)_=0.39, p=0.68) and the dark cycle (Right; Genotype: F_(1,27)_=0.26, p=0.61; State: F_(2,27)_=34.03, p<0.001; Genotype × State: F_(1,10)_=0.042, p=0.96) in WT (n=6) and KO (n=5) VMAT2^DBHcre^ mice. **C**. Average bouts duration in wake, NREM and REM during the light (Top; Genotype: F_(1,27)_=1.33, p=0.26; State: F_(2,27)_=99.72, p<0.001; Genotype × State: F_(2,27)_=0.40, p=0.68) and the dark cycle (Bottom; Genotype: F_(1,27)_=0.27, p=0.61; State: F_(2,27)_=34.03, p<0.001; Genotype × State: F_(2,27)_=0.043, p=0.96) in WT (n=6) and KO (n=5) VMAT2^DBHcre^ mice.

**Figure 7:**
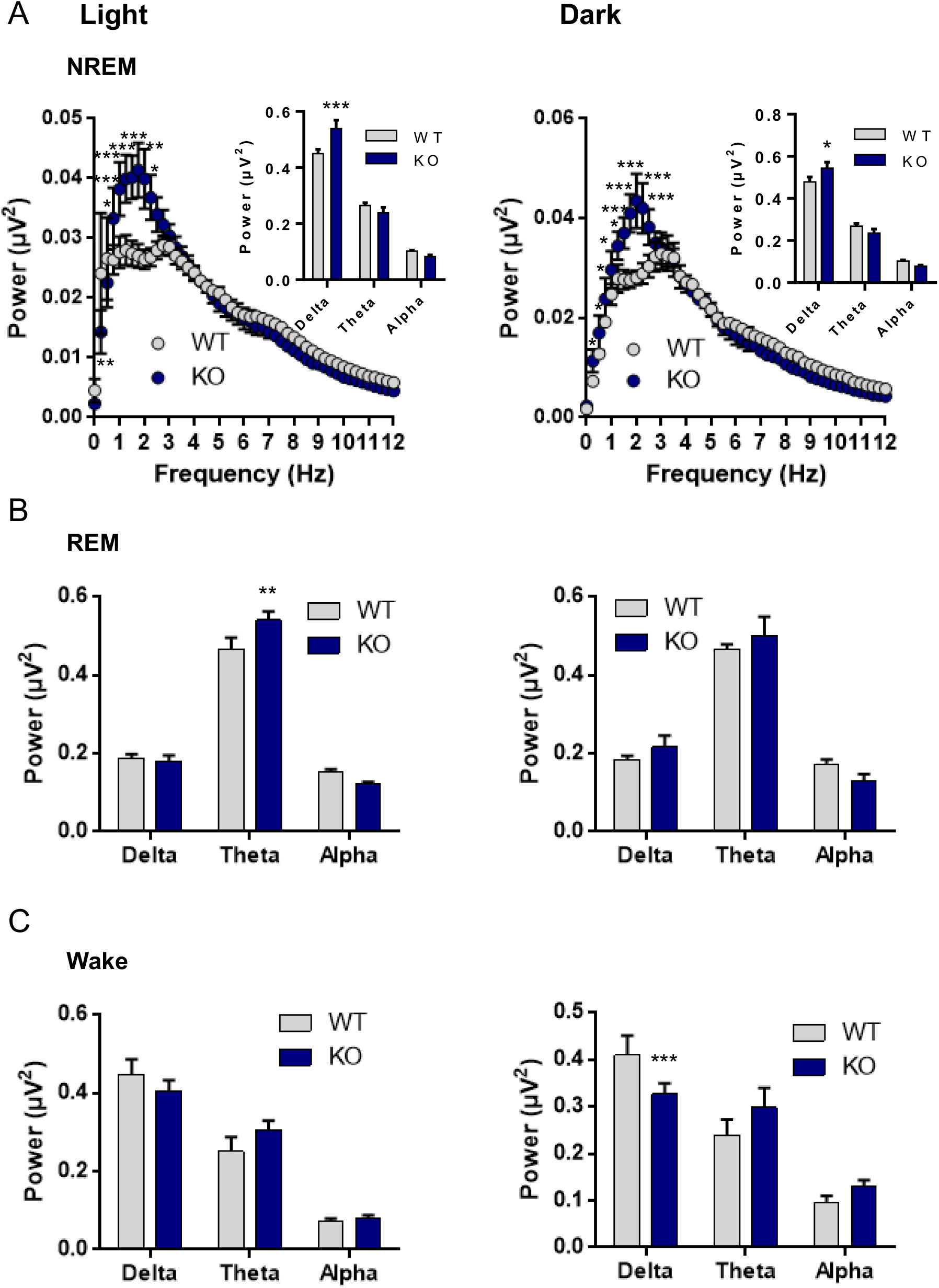
Sleep recording analysis in VMAT2^DBHcre^ mice: power spectrum frequency. **A**. Average spectral distribution of cortical EEG power spectrum during NREM in WT (n=6) and KO (n=4) VMAT2^DBHcre^ mice during the light cycle (Left; Genotype: F_(1,384)_=13.59, p=0.006; Frequency: F_(48,384)_=37.49, p<0.001; Genotype × Frequency: F_(48,384)_=2.50, p<0.001; *post-hoc* test *p< 0.05, **p<0.01, ***p<0.001) and the dark cycle (Right; Genotype: F_(1,384)_=8.15, p=0.021; State : F_(48,384)_=77.11, p<0.001; Genotype × State: F_(48,384)_=3.64, p<0.001; *post-hoc* test *p< 0.05, **p<0.01, ***p<0.001). The power analysis by frequency ranges (Delta: 0.5-4.5 Hz; Theta: 5-9 Hz; Alpha: 9-12 Hz) is shown in the top right corner of each NREM power spectrum distribution curve during the light cycle (Top; Genotype: F_(1,24)_=1.39, p=0.25; State: F_(2,24)_=360.19, p<0.001; Genotype × State: F_(2,24)_=9.31, p=0.001; *post-hoc* test ***p< 0,00032) and the dark cycle (Bottom; Genotype: F_(1,24)_=0.052, p=0.82; State: F_(2,24)_=280.88, p<0.001; Genotype × State: F(2,24)=4.70, p=0.019; *post-hoc* test *p< 0,015). **B**. Wake period power spectrum analysis by frequency ranges (Delta: 0.5-4.5 Hz; Theta: 5-9 Hz; Alpha: 9-12 Hz) in WT (n=6) and KO (n=4) VMAT2^DBHcre^ mice during the light cycle (Left; Genotype: F_(1,24)_=0.091, p=0.76; Frequency: F_(2,24)_=67.79, p<0.001; Genotype × Frequency: F_(2,24)_=1.28, p=0.30) and the dark cycle (Right; Genotype: F_(1,24)_=0.74, p=0.40; Frequency: F_(2,24)_=57.88, p<0.001; Genotype × Frequency: F_(2,24)_=9.20, p=0.0011; *post-hoc* test ***p= 0,00055). **C**. REM period power spectrum analysis by frequency ranges (Delta: 0.5-4.5 Hz; Theta: 5-9 Hz; Alpha: 9-12 Hz) in WT (n=6) and KO (n=4) VMAT2^DBHcre^ mice during the light cycle (Left; Genotype: F_(1,24)_=0.75, p=0.39; Frequency: F_(2,24)_=239.31, p<0.001; Genotype × Frequency: F_(2,24)_=4.37, p=0.024; *post-hoc* test **p=0,086) and the dark cycle (Right; Genotype: F_(1,24)_=0.207, p=0.65; Frequency: F_(2,24)_=144.02, p<0.001; Genotype × Frequency: F_(2,24)_=2.15, p=0.14).

### Transcriptional effect of NE Differential gene expression signature of brain-specific NE depleted mice

#### Profiling gene expression changes across brain regions

To investigate the potential role of NE at the molecular level in maintaining a genetic homeostatic pressure on gene expression, we performed microarray transcriptome profiling in six key cerebral regions including the LC, VTA, Raphe, NAc, PFC and DG of VMAT2^DBHcre^ KO and WT mice.

We first profiled differential gene expression in each brain region of 3-month-old KO versus WT naïve male mice. Table 2 summarizes the number of upregulated (Ups) and downregulated (DNs) differentially expressed genes (DEGs) at different thresholds. When using a low stringency filtering by taking only the adjusted p-value ≤ 0.05 to identify DEGs. The range of DEGs varies from the highest number in the raphe (≈4000) to the smallest number observed in the DG (≈1300). When a more stringent filtering is applied by combining a logFC ≥1, in addition to the adjusted p-value ≤ 0.05, the number of DEGs decreases dramatically, with the DG exhibiting the most DEGs (53, 38 Ups and 15 DNs) while the VTA displays 12 DEGs (8 Ups, 4 DNs).

Figure 8 shows the heatmaps of the DEGs for each region using the most stringent filtering. From this analysis, the SV2c gene with a high potential for neuronal activity regulation was found in four out of the six analyzed regions as the most up-regulated gene in the VMAT2^DBHcre^ KO compared with WT. Hence, this gene was chosen for further validation (see below: probing the biological significance of NE-specific DEGs).

**Figure 8:**
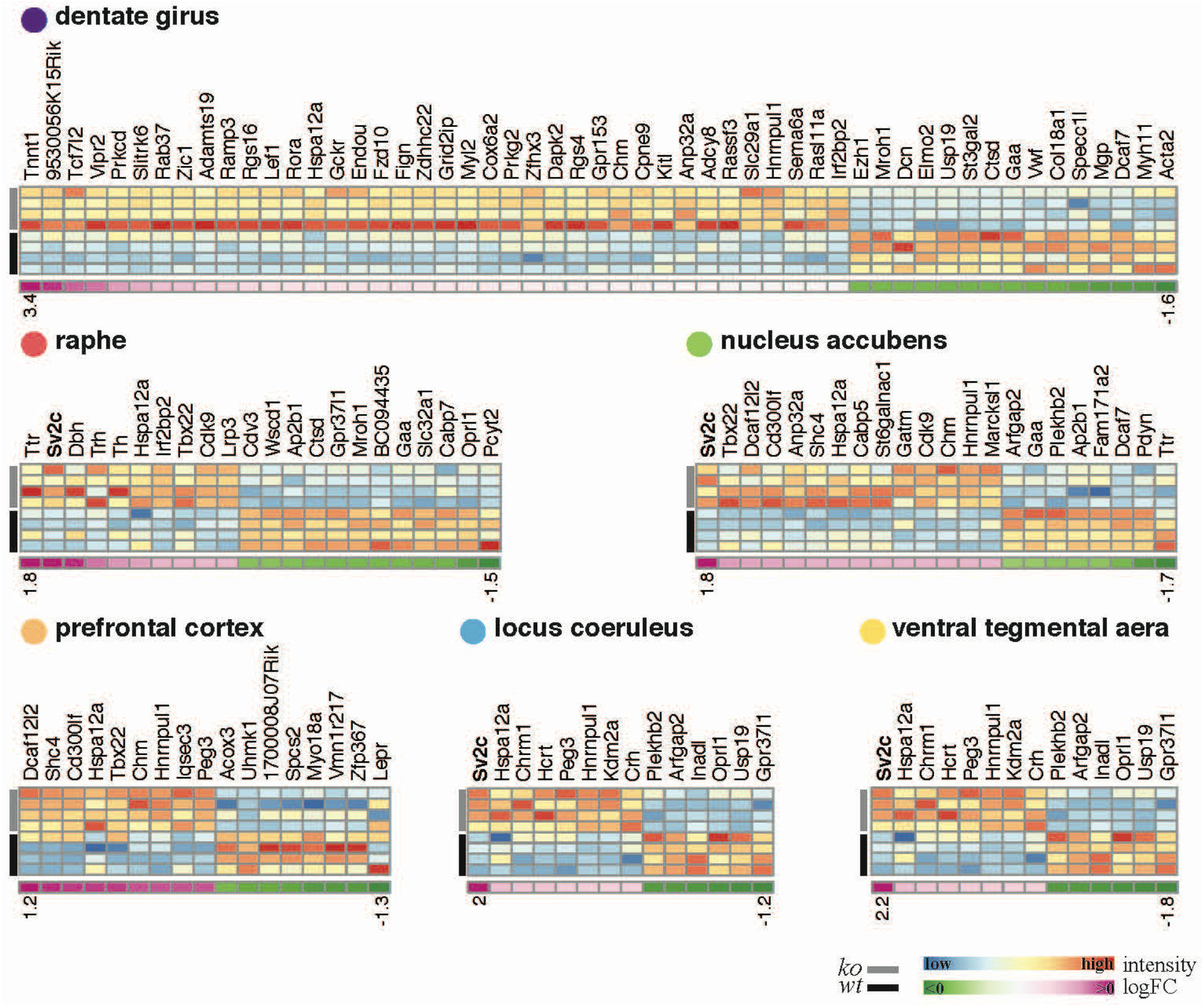
Heatmaps of differentially expressed gene in six brain areas in VMAT2^DBHcre^ mice. Columns represent the differentially expressed genes (i.e adjusted p-value ≥0.05 and |log fold| ≥ 1), with gene symbol on top, and rows represent the samples with the four rows corresponding to WT samples rows color-coded in black and the four rows corresponding to VMAT2^DBHCre^ KO in grey. The color temperature varying from blue (low) to red (high) indicates the intensity of expression. As indication, below each heatmap, a single line heatmap displays log fold information with shades of pink for positive values (i.e genes up-regulated in the KO) and shades of green for negative values (i.e genes down-regulated in KO). The log fold heat map is not scaled with one another and numbers below each extremity (positive and negative) indicates the range of values for each region.

The Gene Set Enrichment Analysis (GSEA) allows using not only the DEGs but all the expressed genes [57], ordered by the t-statistic from the moderate t-test performed, with the most up-regulated genes at the top of the list and the most down-regulated genes at the bottom of the list. This type of analysis has the advantage to be threshold-free, since all the genes are taken into consideration, with the weight of the gene reflected by its position in the ranked list. We used this method to determine whether preexisting gene collections (Hallmark, GO) were affected by NE depletion. Binary heatmaps in Figure 9 show that the effect of the KO has a limited effect on the studied pathways; not all the regions displayed notable alterations and those few affected pathways were related to global metabolism and not neuronal specific (except GO neuropeptide receptor binding, up-regulated in the locus coeruleus).

**Figure 9:**
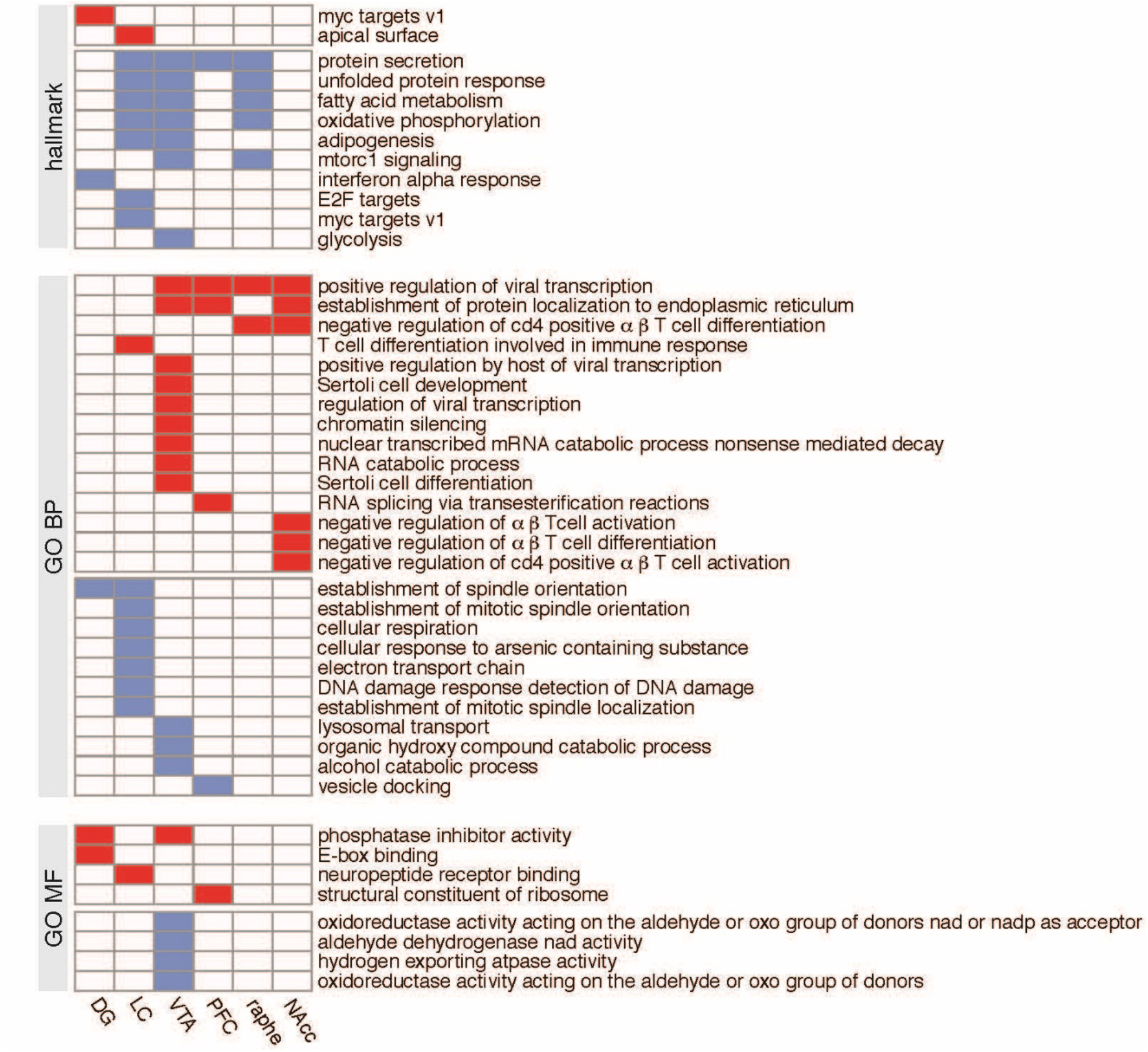
Enriched gene sets across individual brain regions. Enriched (over-represented) gene sets with up-regulated genes in VMAT2^DBHCre^ KO are shown in red; enriched gene sets with down-regulated genes in VMAT2^DBHCre^ KO are shown in blue. Gene set enrichment profiles using the Hallmark or Gene Ontology Biological Process (GO BP), or Gene Ontology Molecular Function (GO MF) gene collections, each row corresponds to a gene set, and each column corresponds to a brain region.

Taken together, these observations indicate a counterintuitively mild effect at the transcriptomic level of the VMAT2^DBHcre^ KO, despite the absence of any paralog that could substitute for VMAT2 deletion. Genetic and cerebral plasticity might have taken place during the development and adolescence phases compensating for the VMAT2 loss in DBH-positive neurons.

#### Probing the biological significance of NE-specific DEGs

The transcriptome mapping used in this study could offer a template to identify NE targets that can be exploited for a better understanding of NE involvement in brain function and behavior. To probe the significance of NE-specific DEGs as novel targets to understand NE signaling function in NE-specific behavioral alteration, we selected the SV2c gene, both because it was the most up-regulated DEG in 3 of the 6 regions sampled (including the LC, VTA, and NAc) and given its previously demonstrated role in neurotransmitters release and action. SV2c radioactive in situ hybridization in the LC, VTA, and NAc of VMAT2^DBHcre^ KO and WT mice allowed us to confirm the increased expression of the SV2c gene in the LC as observed by microarray (t_6_=−4.25, p=0.0054, Figure 10 A-B).

**Figure 10:**
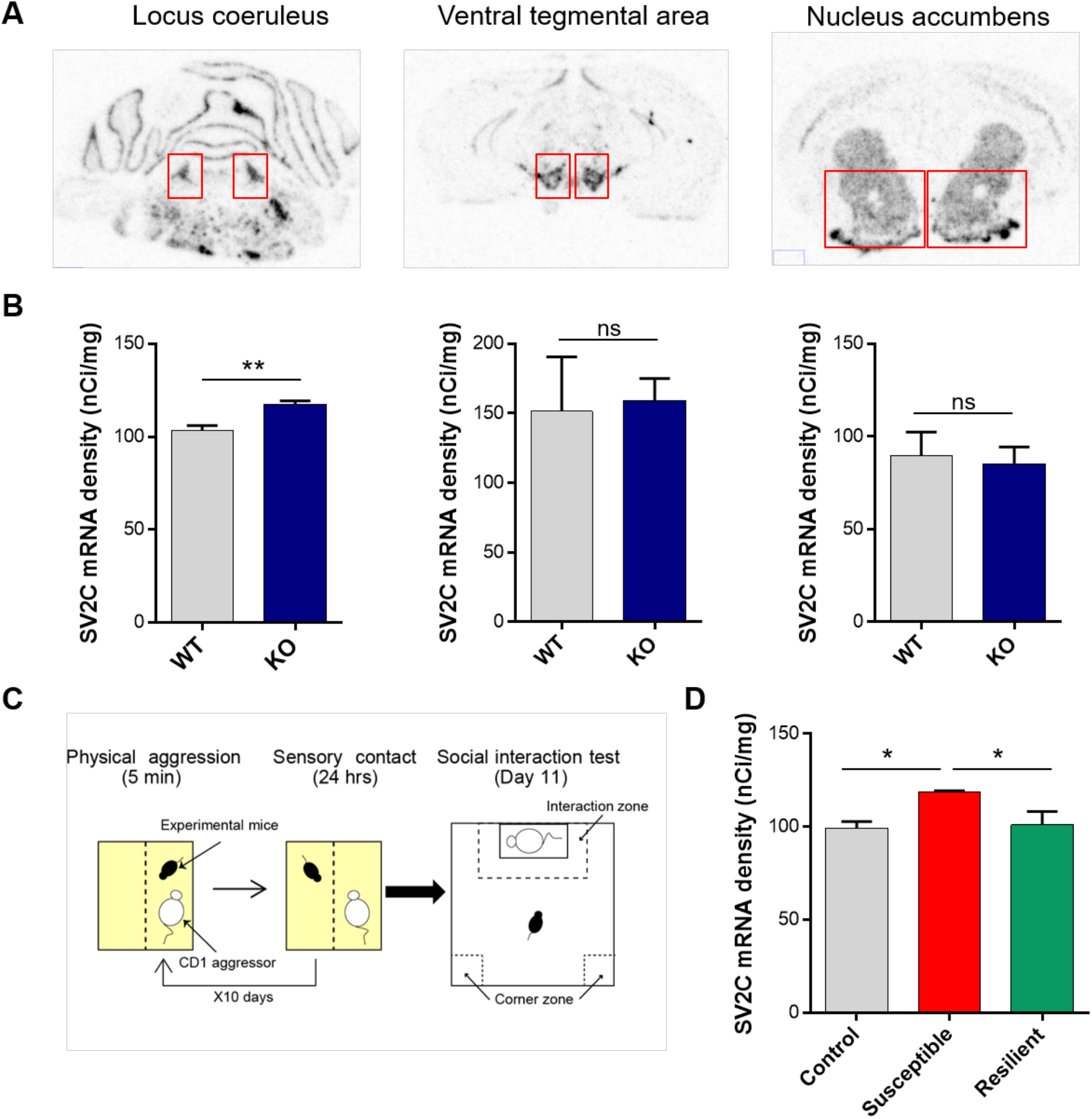
SV2c in situ hybridization. **A**. Illustration of SV2c mRNA radioactive in situ labeling in the locus coeruleus (Left), ventral tegmental area (center), and nucleus accumbens (right). **B.** Density (nCi/mg) of SV2c mRNA expression in the locus coeruleus (Left), ventral tegmental area (center), and nucleus accumbens (right), measured by radioactive in situ hybridization, in WT (n=4) and KO (n=4) VMAT2^BDHcre^ mice (t_6_=−4.25, p=0.0054). **C**. Schematic representation of the chronic social defeat stress paradigm. **D**. Density (nCi/mg) of SV2c mRNA expression in the locus coeruleus, measured by radioactive in situ hybridization, in Control (n=5), susceptible (n=4), and resilient (n=4) mice to the chronic social defeat stress paradigm (F_(2,10)_=5.41, p=0.026; P*ost hoc* test: CTL *vs* SUSC: *p=0.012, SUSC *vs* RES: *p=0.025, CTL *vs* RES: p=0.79, ns).

Based on previous results from our group showing that KO mice with NE-depletion demonstrate a strong susceptible phenotype in response to chronic social defeat stress (CSDS, Figure 10C) [58]. We hypothesized that the expression of SV2c (a putative marker of NE depletion) would be increased in the LC of susceptible mice but not in resilience. We thus looked at SV2c expression by in situ hybridization in the LC of control, susceptible and resilient mice to 10-day CSDS. Remarkably, we found a specific increase in the expression of SV2c in the LC of susceptible mice compared to control and resilient animals (F_(2,10)_=5.41, p=0.026; P*ost hoc* test: CTL vs SUSC: *p=0.012, SUSC vs RES: *p=0.025, CTL vs RES: p=0.79, ns; Figure 10D).

## DISCUSSION

The present study addressed the implication of central NE in regulating/modulating specific behavior and identified the brain networks and molecular mechanisms underlying NE-associated behavioral functions. Table 1 summarizes the overall behavioral results that are discussed below.

**Table 1:** Behavioral results summary in VMAT2^DBHcre^ mice.

| Behavioral analysis |  |  | NE-depletion effects |  |
| --- | --- | --- | --- | --- |
| **Anxiety- and depression-like parameters** | Elevated plus maze |  | ∅ | Anxiolytic and antidepressant effect |
|  | Novelty suppressed feeding test |  | ↘ eating latency |  |
|  | Marble burying |  | ↘ number |  |
|  | Force swim test |  | ↘ immobility |  |
|  | Sucrose preference |  | ∅ |  |
|  | CORT level |  | ↗ CORT level return |  |
|  | Dexamethasone suppression |  | ↗ suppression |  |
| **Analgesia** | Tail immersion |  | ∅ | No analgesic effect |
| **Learning and memory** | Fear conditioning | Context | ↗ immobility | Enhance contextual fear memory |
|  |  | cue | ∅ |  |
|  | Morris water maze | Spatial | ∅ |  |
|  |  | Reversal | ↗ time in target quadrant |  |
|  |  | Rapid place learning | ∅ |  |
| **Executive function** | Attentional set shifting | Reversal | ∅ | Blunt shifting behavior |
|  |  | ID | ∅ |  |
|  |  | ED | ↗ trial number |  |
| **Addiction** | Locomotor drug's response | Cocaine | ∅ | Decrease amphetamine-induced hyperlocomotion |
|  |  | Amphetamine | ↘ locomotor response |  |
|  | Drug's sensitization | Cocaine | ∅ |  |
|  |  | Amphetamine | ∅ |  |
| **Circadian cycle** | Running wheel |  | ↘ activity | Decrease running wheel activity |
|  | Telemetry |  | ∅ |  |
| **Sleep analysis** | % Time | Awake | ∅ | Deeper sleep pressure during NREM |
|  |  | NREM | ∅ |  |
|  |  | REM | ∅ |  |
|  | Boots number and duration | Awake | ∅ |  |
|  |  | NREM | ∅ |  |
|  |  | REM | ∅ |  |
|  | Power spectrum | Awake | ↘ Delta (Dark phase) |  |
|  |  | NREM | ↗ Delta (Dark and light phases) |  |
|  |  | REM | ↗ Theta (Light phase) |  |

**Table 2.**
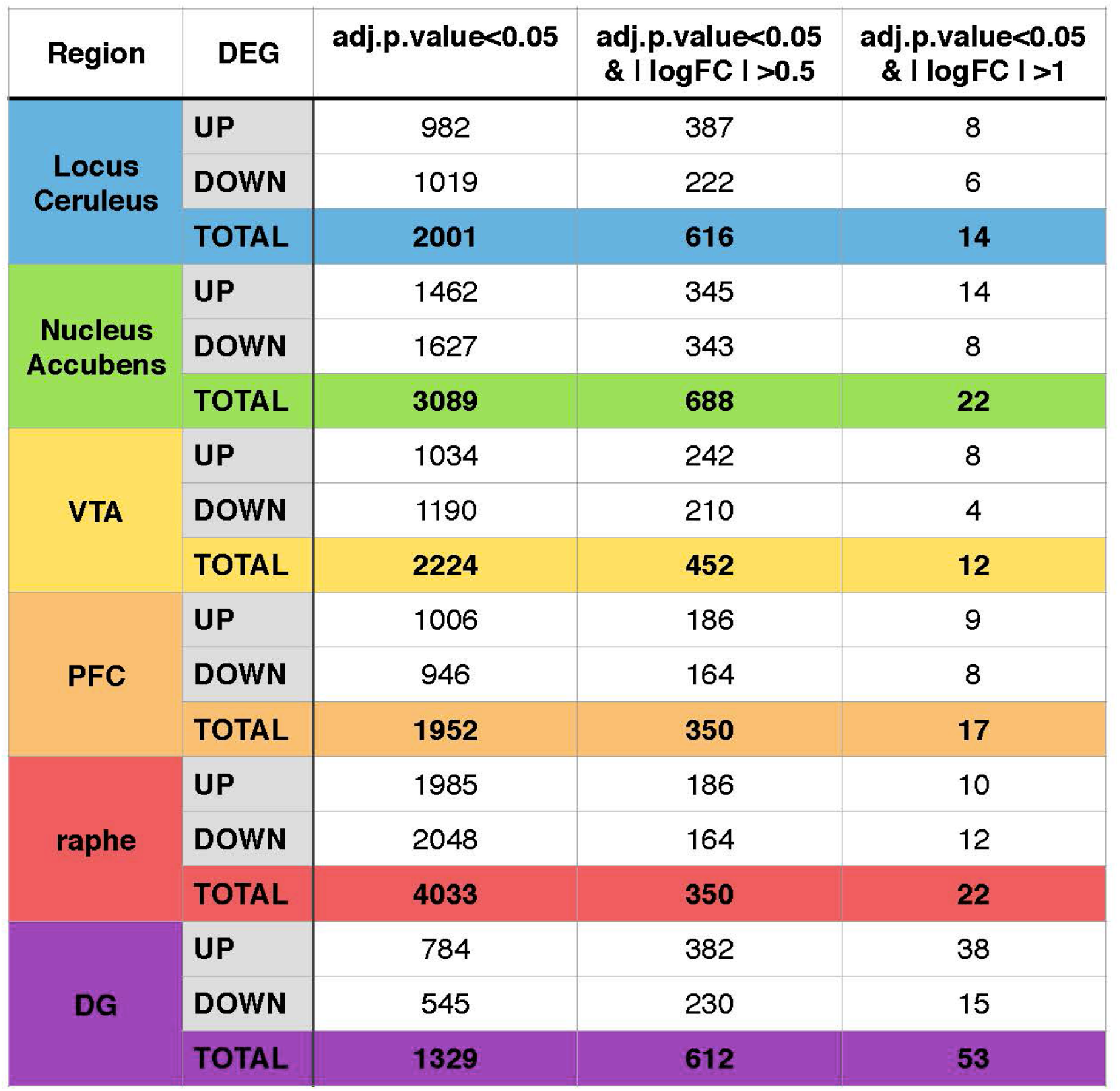
Summary of differentially expressed genes (DEGs) in the different brain regions for different levels of stringency. For each region the number of up-regulated, down-regulated and total DEGs are indicated for different levels of stringency from the less stringent where only the adjusted p-value (≤0.05) is taken into account, to the most stringent where the adjusted p-value ≤0.05 is combined to an absolute value of log-fold (logFC) ≥1.

### Anxiolytic and antidepressant effects of NE depletion

Together with neuronal and hormonal systems, the NE system is activated by stress to induce various physiological and behavioral changes [31,59–61]. The central NE system shares common reciprocal functional connections with both the monoaminergic systems (DA and 5HT) and with the paraventricular nucleus (PVN) of the hypothalamus [62], a major actor of the HPA axis. Thus, dysregulation of the NE system can alter the normal regulation of these systems and precipitate stress-related psychopathologies such as anxiety- and depression-like disorders. Several studies have found dysregulation of the NE system in both anxiety and depressive disorders. For example, anxiety disorders and melancholia are associated with increased NE plasma concentrations [63–65]. Postmortem studies have demonstrated increased NE levels in the brains of unipolar and bipolar suicide victims [66–68]. To investigate whether NE dysregulation in anxiety and depression is a cause or a consequence of the pathology, we investigated the implication of NE in anxiety and depression-like behavior in basal conditions (i.e. not under chronic stress).

Studies looking at NE function in animal models of anxiety/depression-like behavior report heterogeneous findings based to the approach used to manipulate NE synthesis or release. Using either genetic approaches of decreased NE transmission (DBH KO, α1R deletion mice) or pharmacological, neurotoxins and immunotoxin lesions, absence of effect, anxiogenic or anxiolytic effects have been associated with alterations in NE transmission in the EPM, light/dark test, open-field and FST [35,69–76]. Similarly, increasing NE release via blockade of the NET induced antidepressant effect and pharmacological blockade of the α2 autoreceptors produced an anxiogenic behavior [77–80]. Here, we reported anxiolytic and antidepressant-like effects of central NE depletion in the FST, marble-burying test, and novelty-suppressed feeding test (Figure 1A-D). This suggests that an excess of NE without stress exposure can have adverse effects on mood and anxiety. This deleterious effect of excessive NE, proposed as the “noradrenaline paradox” [81], is confirmed by studies reporting that photostimulation of LC-NE neurons projecting to the BLA produced anxiety-like behavior [82]. The absence of effect of NE depletion in the sucrose preference test (Figure 1E) is in line with the observations that tricyclic antidepressant blockade in non-stressed animals does not induce changes in anhedonia [83]. Acute stress is known to rapidly increase circulating adrenaline/noradrenaline, triggering increased corticosterone levels in rodents [61,84]. Interestingly, cortisol responses to physiological stressors in patients with MDD have been found to be significantly higher than for healthy individuals during the recovery period. This suggests that MDD is associated with an inability to turn off the production of cortisol via negative feedback of the HPA axis [85]. In our VMAT2^DBHcre^ KO mice, the blood corticosterone increase following acute stress was similar to WT mice (Figure 1F), because in this model, E and NE releases are fully preserved at the periphery. Very interestingly, the rapid decrease of circulating corticosterone one hour after the end of the acute stress and the increase in dexamethasone-induced corticosterone suppression in VMAT2^DBHcre^ KO (Figure 1F-G) suggested an improvement of the HPA axis negative feedback to control corticosterone release upon NE depletion. These results suggest a direct control of central NE upon its homeostatic regulation at the periphery.

### NE regulation of emotional memory

Controversial studies of NE signaling manipulation demonstrated that reduced NE transmission could either blunt fear acquisition [86–88] or have no effect [89–92]. Regarding fear expression, studies of decreasing NE transmission showed the same discrepancy with a reduction [93] or no effect on cued fear expression [89–92,94]. Whereas these studies used a single shock conditioning trial, it has been shown that NE blockade had no effects on single-trial conditioning, but is necessary for multi-trial enhancement of learning and subsequent expression of that memory [95]. Weaker conditioning protocols are less likely to recruit LC-NE given that they appear insensitive to NE manipulations.

In our study, we demonstrated that fear acquisition was intact in mice lacking NE signaling, however, NE depletion was increasing contextual fear expression without affecting cue fear expression (Figure 2). Interestingly, it has been recently reported that in mice with a global decreased expression of VMAT2 (VMAT2lo), the same increase of contextual fear expression occurs [96]. Therefore, these behavioral changes may be due to VMAT2 decrease in NE neurons. It has been also reported that 6-OHDA lesions of the dorsal noradrenergic bundle (but not the ventral) that produce a very strong decrease of NE projections, also enhanced contextual fear [97,98]. Future studies should be done to test different conditioning protocol intensity in order to identify when the NE system becomes involved in fear acquisition and cued fear expression in VMAT2^DBHcre^ KO mice.

### Absence of NE implication on spatial and working memory

Evidence for a role of NE in memory processes comes mainly from pharmacological studies in which noradrenergic transmission is manipulated after memory acquisition. Pharmacological studies have also revealed a late stage of memory formation that is dependent on β-noradrenergic receptors: rats injected intracerebrally with a β-receptor antagonist 2 hours after learning showed amnesia when tested 48 hours later. If the injection was administered immediately after learning there was no effect, suggesting that there is a time window after a learning experience during which the NE system is activated to reinforce long-term memory processing [99,100]. Many studies using neurotoxic, electrolytic, or pharmacological lesions of the LC have found effects on various aspects of learning and memory including fear learning, extinction and reversal learning, avoidance, and working memory [101–108]. However, this role of LC neurons in many different types of learning and memory has not been replicated consistently [98,102,109–111]. This variability can be explained by the type and size of the lesion and the behavioral outcome but also by compensatory mechanisms.

Remarkably, we demonstrated that NE depletion does not induce learning and memory deficit since performances of mice lacking NE signaling are equal and even better (or faster) compared to control mice in multiple versions of the Morris water maze test (spatial version, reversal, and working memory, Figure 3A-C). Because in the VMAT2^DBHcre^ KO mice, NE neurons do not degenerate following lesions but rather unable to release DA as it has been recently suggested [112,113], it remains possible that other neurotransmitters present in LC-NE neurons, including Galanin or NPY for example [114], could balance NE loss in the memory performances.

### NE depletion in executive functions and behavioral flexibility

Behavioral (or cognitive) flexibility is a brain mechanism allowing one individual to adapt to a changing environment. We used the two most classical rodent tests to evaluate behavioral flexibility, a reversal learning test using the Morris water maze and the Attentional Set-Shifting Test (ASST) which is a rodent adaptation [115] of the Wisconsin Card Sorting Test in human. Whereas 5HT and DA transmission are clearly implicated in behavioral flexibility mechanisms [116,117], a role for noradrenergic transmission has been also proposed [118]. In the reversal learning test, the VMAT2^DBHcre^ KO mice showed better performance, they can shift their learning for a new platform location more efficiently than WT mice (Figure 3C). Even though this observation seems to contradict the identified role of NE projection to the frontal cortex in the reinforcing learning mechanisms [13], the disappearance of NE transmission could favor the role of DA in this process [119]. In the ASST, mice learned to retrieve a reward by focusing on a sole perceptual feature of a complex stimulus. KO mice were able to perform the task as well as WT mice in the simpler trials of the task (i.e., when the reward was associated with only an odor, or only a medium). However, when the difficulty of the task increased (extra-dimensional shift), the KO mice showed a deficit in performance as they were not able to unlearn the previous association and learn a new one (Figure 3D). This highlights the functional role of NE transmission in task shifting behaviors.

### Role of NE depletion in locomotor response and behavioral sensitization to psychostimulants

Psychostimulant drugs are directly targeting the three aminergic transporters, DAT, SERT and NET, with an overall effect of locomotor increase and addictive properties [6,120]. Even if the DAT is the main target of cocaine and amphetamine [121,122], NE increase by NET inhibition plays a functional role [123,124]. The acute effect of cocaine on locomotion were the same between WT and VMAT2^DBHcre^ KO mice, showing the main role of DAT blockade in that case (Figure 4A). For amphetamine, interestingly we observed a lesser effect for all doses on locomotion (Figure 4C). Amphetamine, in addition to its transporter inhibition properties, also directly targets VMAT2 and is responsible for a non-exocytic release of amines [125]. Therefore, NE releases following amphetamine administration is likely partially responsible for the locomotor increase observed in WT mice, as it has been also observed in the NETKO mice in which VMAT2 could still be targeted by amphetamine [124] and where amphetamine has significant locomotor effects.

Interestingly, we observed no differences in behavioral sensitization to cocaine or amphetamine after one week of chronic administration (Figure 4B,D). This seems to rule out a role for NE transmission in this process, as has been earlier suggested using the a1-adrenergic antagonist prazosin administration in the PFC upon amphetamine administration [126], but not replicated in self-administration studies [127]. Besides the explanation of developmental adaptation in our model, another possible reason for the lack of effect could be that α1-adrenergic receptors are co-localized with D1 dopamine receptors [128], and therefore the effect of prazosin could likely occur even in the absence of NE release, a hypothesis that remains to be investigated in the VMAT2^DBHcre^ KO mice.

### Circadian rhythms

24-hour rhythms in physiology and behavior are governed by the master circadian clock which resides in the suprachiasmatic nuclei (SCN) of the hypothalamus, but other clock gene-expressing brain regions may contribute as well [129,130].

Accumulating evidence suggests that the clock system is interconnected with the HPA axis via synaptic connections between the SCN and the PVN [131–133].

Interestingly, the stress system, through the HPA axis, communicates with the clock system; therefore, any uncoupling or dysregulation could potentially cause several disorders, such as metabolic, autoimmune, and mood disorders [134]. Disruption of the physiological circadian rhythms induces an increase in NE level both in the LC and in the PVN [135]. A meta-analysis looking at the level of NE in different projection areas during the light/dark cycle highlighted a higher NE level during the dark phase and a stable or lower level during the light phase. Some studies reported NE to peak between 1 and 3 hrs after the dark period onset, especially in the PVN. Sleep deprivation induces an increase NE level during the deprivation in the medial PFC or an increase after the deprivation in the NAc [136].

We did not find differences in locomotor activity rhythm periods between WT and VMAT2^DBHcre^ KO animals in constant darkness indicating that the NE deficit does not affect circadian rhythm generation and thus SCN clock function (Figure 5A). We also found total daily home cage activity derived from telemetry implant recordings to be normal, however, VMAT2^DBHcre^ KOs showed lower daily totals in running wheel activity, which could reflect altered arousal regulation (Figure 5C-F).

### NE and sleep and waves

LC-NE is implicated in arousal, as suggested by the original finding that LC neurons are active during wake, decrease firing rate during NREM sleep, and are silent during REM sleep [137]. Depletion of the dopamine-ß-hydroxylase (DBHKO mice) generates mice with altered sleep and arousal patterns who tend to sleep more overall, but with less REM sleep over a 24-hour period [138,139]. Of importance, DBHKO mice have a total absence of NE and E in the whole body [43] and most of them die early, due to the absence of peripheric NE and E. Chemical rescue of DBHKO mice using L-threo-3,4-dihydroxyphenylserine (DOPS) in the maternal drinking water did not rescue the survival rate of the animals (30 to 40%) nor did it restore NE and E in all organs including several brain regions and the adrenal glands [140]. Using CRISPR/Cas9 technology to disrupt the DBH gene selectively in adult LC-NE neurons, NE depleted mice have reduced wake length even in the presence of salient stimuli and increased NREM sleep amount [141]. This study is in accordance with our data which suggest that the lack of NE release led to a decrease in arousal concomitant with an increase of sleep (only during some nocturnal hours), a sleep feature that becomes more prevalent on the power spectrum analysis (Figure 6). Indeed, the slow wave activities in the cortical EEG power spectrum during NREM were significantly increased in VMAT2^DBHcre^ KO mice as compared to controls (Figure 7). This increase was similar to what could be observed in WT animals with optogenetic inhibition of neuronal activity in the LC [14]. Collectively these data indicate that the absence of NE release led to changes in sleep quality rather than sleep quantity.

### Transcriptomic analyses

We used microarray technology to identify transcriptional changes occurring in the absence of CNS NE, therefore genes that are likely under regulatory control of NE transmission. In addition to the LC which comprises most of the NE neurons in the brain, we also investigate brain regions formerly identified for receiving NE input [142–144]. These regions implicated in cognitive and emotional functions, include the VTA, raphe nucleus, DG of the hippocampus, NAc, and PFC. At the higher stringency threshold (log2 fold change > 1; P<0.05), the total number of differentially expressed genes (DEGs) revealed by RNA microarray was not very high, with the lowest number of DEGs in the LC and the VTA (14 and 12, respectively) and the highest number of DEGs in the dentate gyrus (54) (Table 2). The fact that the LC appeared to be the least affected region was a striking finding that could be associated with the absence of a “major”phenotype of the VMAT2^DBHcre^ KO mice, as compared to the phenotypes of the VMAT2^DATcre^ KO mice and VMAT2^SERTcre^ KO mice [47,145–147]. This absence of NE role during brain early development is also in agreement with observation in humans with noradrenaline deficiency [148] or mice with a constitutive deletion of DBH [43], of a cardiac, but not central, phenotype.

We also conducted a gene set enrichment analysis (GSEA) using the Hallmark, GO molecular function, and GO biological process gene collections (Figure 9). Interestingly, in the LC we mostly identified increased expression for several genes, including those implicated in spindle organization or protein secretion, potentially linked to changes in vesicular organization associated with the absence of VMAT in LC-NE neurons. Of particular interest is the finding that the largest number of up- or down regulated genes observed in GO classes was in the VTA, which could be a consequence of the direct inhibitory role of NE on DA-VTA neurons as previously reported [58].

The heatmap representation of all individual up- or down-regulated genes revealed the striking finding that for 4 of the 6 brain regions analyzed (LC, NAc, VTA, and raphe nucleus), the same gene, SV2c, is the most highly up-regulated gene (Figure 8). We therefore decided to validate the functional relevance of our differential transcriptomic analysis by further functional investigation with the SV2c gene.

### SV2c expression is regulated by NE transmission and chronic stress

The Synaptic Vesicle Glycoprotein 2c (SV2c) is a glycoprotein with twelve transmembrane domains found on vesicular membranes [149], and highly related to SV2a and SV2b. While their functional roles are not yet clearly identified, SV2 proteins were found to be the receptors for the Botulinum neurotoxin neuronal entry [150]. In contrast to SV2a and SV2b, SV2c has a restricted localization in the brain, with no expression in the cortex and the hippocampus [149,151,152], the two brain regions in our transcriptional analysis that did not show any increased expression. Genetic deletion of SV2c has been shown to profoundly affect DA neuron dynamics [153,154]. SV2c has been associated with the DA system in previous research. For example, a correlated decrease of SV2c was found in the post-mortem brain of Parkinson patients [154], whereas in a 6OHDA mouse model, a DA lesion is responsible for an increased expression of SV2c[153]; the difference in these observations between animal models and human pathology could have several explanations, though it remains strongly indicative of SV2c plasticity in these DA neurons. However, so far, its role has not been investigated in the NE system.

First, we confirmed SV2c increased expression in VMAT2^DBHcre^ KO mice using quantitative in situ hybridization with radiolabeled probes (Figure 10A-B). Only in the LC, we were able to verify a significant 20% increase of SV2c mRNA levels in the VMAT2^DBHcre^ KO mice. Previous work established that chronic social defeat stress (CSDS) induces two phenotypes: mice that are susceptible to stress (~70%) and exhibit social avoidance, and those that are resilient against stress (~30%) and continue to show a preference for social interaction like control mice [155]. In VMAT2^DBHcre^ KO mice with brain-specific NE depletion, we observed a profound increase proportion of susceptible animals in response to chronic social defeat, KO mice have a strong susceptible phenotype [58]. We were able to confirm our hypothesis that an increase in SV2c expression could be a marker of NE-depletion induced susceptibility to stress by demonstrating that there was a 20% increase in SV2c expression in the LC of susceptible mice compared to control mice, but not in resilient mice (Figure 10C-D). Although the presynaptic mechanisms responsible for NE release alteration in stress susceptibility are unknown, the increased SV2c expression observed in the LC of susceptible mice could either be the presynaptic mechanisms responsible for the NE release dysregulation or serve as a compensatory mechanism aimed at normalizing NE release in this condition. This functional correlation fully validates the differential transcriptomic analysis that was conducted and could offer interesting targets for further validation.

## CONCLUSION

Despite the brain-wide distribution of NE that makes it a challenging system to target, the NE system remains a viable target for new drug discovery. A better understanding of NE influences in cognitive control and executive function could offer therapeutic opportunities for disorders such as attention-deficit hyperactivity and the cognitive deficits observed in schizophrenia, bipolar disorders, and dementia. Moreover, unraveling the exact implication of NE in anxiety, mood, and vulnerability to stress could help ameliorate therapeutic strategies for anxiety and depressive disorders, especially when only 30% of depressed patients achieve complete remission after a single antidepressant trial. In VMAT2^DBHcre^ KO mice with an unprecedented absence of central NE release, there is no strong phenotype, neither developmental nor behavioral, even though NE play a role in several brain functions. The NE system appears to be quite silent at the constitutive level, only when the system is challenged, by either acute or chronic stress, do differences emerge due to central NE depletion pointing to its critical role in stress modulation.

## Author Contributions

Conceptualization, BG with EI; Methodology and investigation for behavioral experiments, EI with CG, LP, EG and VG; Methodology and investigation for circadian rhythms, IDB and KFS; Methodology and investigation for sleep, JCM and AA; Methodology and investigation for transcriptomic analysis, SM; writing—review and editing, BG and EI with MB, EG, SM, AA and KFS; All authors have read and agreed to the published version of the manuscript.”

## Funding

This work was supported by grants from the Canadian Institute for Health Research (201309OG-312343-PT; 201803PJT-399980-PT) and the Graham Boeckh Foundation to BG, the Natural Sciences and Engineering Research Council of Canada (RGPIN-2015-04034) to KFS.

## Data Availability Statement

All data are publicly available. Request for mice shoulc be directed to BG.

## Acknowledgments

We thank Erika Vigneault and Yeqing Geng for taking care of the mice colonies. BG is a Canada Research Chair and a visiting professor at Université de Paris.

## Competing interests

The authors declare no competing interests.

## Notes

### Competing Interest Statement

The authors have declared no competing interest.

